# Immunobiochemical reconstruction of influenza lung infection - Melanoma skin cancer interactions

**DOI:** 10.1101/331546

**Authors:** Evgeni V. Nikolaev, Andrew Zloza, Eduardo D. Sontag

**Affiliations:** Center for Quantitative Biology, Rutgers University, Piscataway, NJ 08854, USA; Section of Surgical Oncology Research, Division of Surgical Oncology, Rutgers Cancer Institute of New Jersey, New Brunswick, NJ 08903, USA; Department of Surgery, Rutgers Robert Wood Johnson Medical School, New Brunswick, NJ 08903, USA; Department of Electrical and Computer Engineering, and Department of Bioengineering, Northeastern University, Boston, MA 02115, USA; Laboratory for Systems Pharmacology, Harvard Medical School, Boston, MA 02115, USA

## Abstract

Our recent experimental results that combine a mouse model of influenza A virus (IAV) infection (A/H1N1/PR8) and a highly aggressive model of infection-unrelated cancer, B16-F10 skin melanoma, showed that acute influenza infection of the lung promotes distal melanoma growth in the dermis of the flank and leads to decreased host survival. Here, we proceed to ground the experimental observations in a mechanistic immunobiochemical model that incorporates the T cell receptor signaling pathway, various transcription factors, and a gene regulatory network (GRN). A core component of our model is a biochemical motif, which we call a Triple Incoherent Feed-Forward Loop (TIFFL), and which reflects known interactions between IRF4, Blimp-1, and Bcl-6. The different activity levels of the TIFFL components, as a function of the cognate antigen levels and the given inflammation context, manifest themselves in phenotypically distinct outcomes. Specifically, both the TIFFL reconstruction and quantitative estimates obtained from the model allowed us to formulate a hypothesis that it is the loss of the fundamental TIFFL-induced adaptation of the expression of PD-1 receptors on anti-melanoma CD8+ T cells that constitutes the essence of the previously unrecognized immunologic factor that promotes the experimentally observed distal tumor growth in the presence of acute non-ocogenic infection. We therefore hope that this work can further highlight the importance of adaptive mechanisms by which immune functions contribute to the balance between self and non-self immune tolerance, adaptive resistance, and the strength of TCR-induced activation, thus contributing to the understanding of a broader complexity of fundamental interactions between pathogens and tumors.

## 1 Introduction

Our experimental studies (Kohlhapp et al., 2016) with a mouse model of influenza A virus (IAV) infection (A/H1N1/PR8) and a highly aggressive model of infection-unrelated cancer, B16-F10 melanoma, demonstrated that melanoma-specific CD8+ T cells are shunted to the lung in the presence of the infection, where they express high levels of an activation / exhaustion receptor, PD-1. We also showed that acute influenza infection of the lung promotes distal melanoma growth in the dermis of the flank and leads to decreased host survival.

These published data (Kohlhapp et al., 2016) demonstrate that infection-induced changes in anti-tumor responses are independent of 1) the tumor type, 2) the site of tumor challenge, 3) the mouse strain, 4) the infectious agent, 5) the site of infection, 6) the use of transplantable tumors, 7) general suppression of viral clearance in the context of a tumor, 8) not due to immune system inability to respond to concomitant challenges, and 9) reversed by PD-1 blockade.

Thus, these findings (Kohlhapp et al., 2016) have revealed a previously unrecognized factor, acute non-oncogenic infection, in promoting cancer growth, pointing out to another important immunological axis in a decision-making process utilized by the immune system when faced with concomitant challenges.

The analysis of the new phenomena, using both immunobiochemical reaction network reconstruction and mathematical modeling, constitutes the subject of our work with the objective to get valuable insight into complex experimental data (Kohlhapp et al., 2016).

To explore and explain these experimental data, as well as other relevant observations known from the literature, phenotypic modeling has been used as an indispensable part of the work. Indeed, the integration of the reconstruction and the phenotypic modeling turned out to be an essential ingredient of our research for the following reasons.

First of all, we note that our immunobiochemical reconstruction of a core reaction network relevant to the experimental observations (Kohlhapp et al., 2016), has been recognized as belonging to the class of Incoherent Feed-Forward Loops (IFFLs) describing a fundamental law of adaptation (Alon, 2006). In our case, this corresponds to the adaptation of the expression of PD-1 receptors with respect to increasing amounts of antigen (Ag), when the level of PD-1 receptors first sharply increases and then decreases.

We then put forward a hypothesis that it is the loss of Ag dose-dependent adaptation of the expression of PD-1 receptors in the anti-tumor CD8+ T cells that is a major factor resulting in multiple effects in the presence of the acute non-oncogenic infection, including: (1) shunting antitumor CD8+ T cells to the inflamed lung, (2) sequestration of the relocated anti-tumor CD8+ T cells in the lung, and (3) related to (1) and (2) accelerated tumor growth.

Therefore, we suggest that the formulated hypothesis constitutes the essence of the previously unrecognized immunologic factor.

To better understand the relevance of the hypothesis to explain the host immune response to self (tumor) and non-self (infection) antigens, we needed a quantitative estimate for Ag ranges for which complete adaptation could be possible.

In order to obtain such a quantitative estimate, we formulated a coarse-grained mathematical model which was then used to obtain the estimate. The estimate turned out to be approximately equal to at least a 10^3^-fold increase in cognate Ag levels counted from the onset of the sharp increase in PD-1 numbers, required to complete the adaptation.

To this end, it was reasonably to assume that viruses replicating with an exponential rate could produce rapidly very large amounts of Ag, sufficient to complete the PD-1 expression adaptation during very short periods of time when acute infection first develops and then is cleared by the immune system. Such periods of times usually last 7 - 10 days.

Because tumor cells cannot replicate as fast as viruses do, it was then reasonable to assume that growing and replicating tumor cells were unlikely to produce enough antigen to complete the PD-1 expression adaptation during similarly short time intervals. In this work, we demonstrate how these arguments help shed light into experiments obtained in (Kohlhapp et al., 2016).

Before we proceed with the description of our results, a few words should be said about the selected quantitative biology and phenotypic modeling approach. Indeed, because biological networks and, more broadly, biological phenomena are manifestly complicated, the phenotypic modeling approach is often used as an alternative to large-scale mechanistic modeling (Hopfield, 1974; McKeithan, 1995; Sontag, 2001; Heinrich and Rapoport, 2005; Warmflash and Dinner, 2009; Sciammas et al., 2011; Martinez et al., 2012; Lever et al., 2014; Galvez et al., 2016; Sontag, 2017). To this end, the phenotypic modeling approach is well-characterized and recognized as “simple enough to be studied mathematically, but not oversimplified to the point of losing contact with the experimental data” as insightfully argued by Gunawardena (2014). A the same time, large-scale mechanistic or semi-mechanistic models construction is usually hampered by the general issue of sparse experimental measurements (De Boer and Perelson, 2013; Eftimie et al., 2016).

In our modeling studies, we also follow Eftimie et al. (2016) who distinguish two general purposes for mathematical models: (*i*) to offer some general theoretical understanding for a biological problem, which does not require any formal model validation, and (*ii*) to help make predictions, which should already depend on a formal model validation.

Throughout this work, whenever we refer to “modeling studies” we actually mean the purpose (*i*) in which case our modeling goal is to develop a phenotypic model to be partially confirmed by showing agreement between observation and simulation.

Our work is organized as follows. Sect. 2 describes computational approaches used. Sects. 3.1-3.3 provide a detailed bibliomics-based reconstruction of mechanisms that explain the influenza-tumor interactions observed in (Kohlhapp et al., 2016). Here, the molecular basis used to formulate our core Triple Incoherent Feedforward Loop (TIFFL) is discussed, and the TIFFL is presented. In Sect. 3.4.2, the TIFFL-based pathway is customized to address immunobiochemical processes occurring under different inflammation conditions in the infected lung and tumor site. Modeling of the TIFFL is described in Sect. 3.5. In Sect. 4, we discuss a modeling outlook to include more details relevant to T cell trafficking phenomena (Sect. 4.1). We conclude the discussion section by formulating a plausible hypothesis suggesting a concise insight into the “previously unrecognized immunological factor” and also provide a more detailed justification of modeling efforts implemented in this work. Finally, the discussion of reactivation of exhausted effector T cells, technical aspects of our mathematical model derivation, computational approaches, including Global Sensitivity Analysis (GSA), and additional figures can be found in the Supplemental Information (SI).

## 2 Materials and methods

The steady-state solutions of the models developed in this work, the solution stability (Sontag, 1998), as well as the parameter continuation of the steady-state solutions (Kuznetsov, 2013) have been studied numerically (Khibnik et al., 1993), using the command-line functionality of matcont6p10, a Matlab^®^-based Continuation Toolbox (Dhooge et al., 2008).

To carry out Global Sensitivity Analysis (GSA), GSAT, a Matlab^®^-based Global Sensitivity Analysis Toolbox was used (Cannavó, 2012). In all cases, we also used Sobol’s quasi-random distribution (Saltelli et al., 2008; Cannavó, 2012) in order to sample parameter values on the corresponding logarithmic scales with the sample size *N* = 2.0 × 10^4^ (Sect. SI-6).

Matlab^®^ Parallel Computing Toolbox was employed whenever possible.

Finally, the color maps were generated using `varycolor.m`, a Matlab^®^-based function.

## 3 Results

A key challenge in the study of T cells within different dual immunological self and non-self contexts, is the organization of large amounts of relevant molecular and biochemical information, which we discuss below in terms of our immunobiochemical reconstruction based on the available bibliomics.

Our primary goal here is to address specifically the ability of T cells to discriminate between infected and tumor cells, presenting non-self and self antigens, respectively, based on differences in pMHC affinity, encoded by significant differences in the values of *k*_off_, the off-rate constant for the antigen (Ag) and the TCR bond, as defined below, which force T cells to respond differently to Ags of both varying affinity and dose.

### 3.1 Immunobiochemical reconstruction scope

To provide quantitative mechanistic insight into experimental observations (Kohlhapp et al., 2016), we first discuss relevant immunological and molecular details available from the literature in order to obtain a more formal immunologic reconstruction relevant to the observations. We then delineate key genetic and signaling pathways, including critical elements of their complex regulation.

Specifically, we will formulate hypotheses and mechanisms to shed light on the following key observations reported in (Kohlhapp et al., 2016):

**(O1)** *Distant* influenza-melanoma interaction: Influenza-induced loss of anti-melanoma CD8+ T cells from the tumor micro-environment (TME) and their sequestration in the infected lung.
**(O2)** The host immune system’s *ability* to respond to concomitant infection challenges: IAV infection does *not* impede clearance of vaccinia virus (VACV) infection under the same conditions, *nor* tumor challenge changes the ability of the immune system to eliminate the infection.
**(O3)** *Reactivation* of exhausted (T_EX_) anti-melanoma CD8+ T cells after anti-PD-1 (*α*PD-1) blockade: (*i*) reactivated anti-melanoma CD8+ T cells which continue to reside in the TME gain thier ability to contribute to the anti-tumor immune response and, additionally, (*ii*) reactivated anti-melanoma CD8+ T cells sequestered in the infected lung may regain their motility and return back to the TME, where they also aid in the anti-tumor response. While primarily focusing on observations **(O1)** - **(O3)**, we will also provide insight into additional important observations in (Kohlhapp et al., 2016):
**(O4)** Infection early in tumor formation *reduces* host survival by promoting tumor growth in the infected host.
**(O5)** *Differential* expression of PD-1 receptors by effector cells (T_EFF_) in the infected lung: Anti-melanoma CD8+ T cells express larger amounts of PD-1 receptors than anti-influenza CD8+ T cells under the same conditions in the infected lung.

Given the complexity of the processes relevant to the observations **(O1)** - **(O5)**, we must limit the scope of our research. Because the interaction between PD-1 and PD-L1 is regarded as a major “T cell brake,” and because it is also a central topic in (Kohlhapp et al., 2016), we center our analysis on the construction of the gene regulatory networks (GRNs) directly involved in the expression of PD-1 receptors under different immunologic contexts. We use a coarse-grain representation of signal transaction networks leading to the initiation or repression of the GNRs considered.

A very important topic that shapes the scope of our work is that T cell activities are controlled at multiple levels. These regulatory controls are necessary to prevent T cells from becoming hyperactivated, causing significant collateral damage to non-target tissue. These types of responses enhance inflammation, resulting in the release of self-antigens from necrotic tissue, increasing the chances for the induction of autoimmune diseases (Liechtenstein et al., 2012). To avoid autoimmunity induced by necrotic tissue, key regulatory T cell inhibitory interactions occur between PD-L1 expressed on immune, infected and tumor cells, and, PD-1 expressed on T cells (Sakaguchi et al., 2008; Fife et al., 2009; Francisco et al., 2010; Schietinger and Greenberg, 2014; Bardhan et al., 2016; Sharpe and Pauken, 2017).

### 3.2 A brief TCR activation primer

For a strong CD8+ T cell activation, three well-known signals have to be provided from professional antigen presenting cells (APCs) (Kindt et al., 2007):

- Signal 1: Antigen presentation to T cells.
- Signal 2: Co-stimulation.
- Signal 3: Cytokine priming.

Signal 1 is mediated by binding of a T cell receptor (TCR) on T cells with its cognate antigen presented on an MHC.

Signal 2 is mediated by a series of receptor:ligand bindings, such as CD80 binding to CD28 between the APC and the T cell, respectively. The combination of TCR engagement, CD28 binding, and IL-2 activates Zap-70, lck and PI3K, which in turn lead to T cell activation, expansion, and acquisition of effector activities (Liechtenstein et al., 2012; Rendall and Sontag, 2017). Furthermore, in reality, as Liechtenstein et al. (2012) states, a variety of ligand-receptor interactions take place in the immunological synapse, many of which are inhibitory. The final integration between activatory and inhibitory interactions determine the type and strength of the co-stimulatory signal given to the T cells, setting the “degree” of T cell activation.

Signal 3 is mediated by binding of cytokines to their respective receptors, such as IL-2 produced by T cells binding to IL-2 receptors also on the same T cells.

During antigen presentation to naï ve T cells, PD-1:PD-L1 interaction acts as a brake in TCR signal transduction. PD-1 is transiently up-regulated during antigen presentation as a consequence of T cell activation (Freeman et al., 2000) and PD-1:PD-L1 binding results in ligand-induced TCR down-modulation (Escors et al., 2011; Karwacz et al., 2011, 2012).

To this end, Liechtenstein et al. (2012) suggest that TCR down-modulation is absolutely required for T cell activation in order to prevent T cell hyperactivation by terminating TCR signal transduction. In such cases, PD-1 associates to the TCR at the immunological synapse and controls its signal transduction as well as its presence on the T cell surface (Karwacz et al., 2011). TCR down-modulation is largely reduced when PD-L1 is silenced in antigen-presenting DCs, or when PD-1:PD-L1 is blocked using antibodies during antigen presentation (Liechtenstein et al., 2012).

Finally, as further reviewed in (Liechtenstein et al., 2012), PD-1:PD-L1 interactions control the timing of TCR stimulation in at least two different ways: (*i*) by removing TCRs from the T cell surface, and (*ii*) by terminating the intracellular signal transduction pathways after recruiting phosphatases SHP1 and SHP2 (Zhang and Rundell, 2006; Bardhan et al., 2016). Note briefly that processes in (*ii*), put, for example, a brake on NF-*κ*B signaling, the inhibitory process that shuts down IRF4, which in turn removes the Blimp-1 imposed brake from PD-1 transcription, where both IRF4 and Blimp-1 are key molecular species of our analysis, as discussed below in detail.

### 3.3 Linking observations with immunological mechanisms

Our bibliomic-wide immunologic reconstruction of infection-tumor interactions (Kohlhapp et al., 2016) is summarized in Table 1.

**Table 1:** A summary of the immunological reconstruction of infection-tumor interactions^a^.

| Observation | Description | Mechanisms |
| --- | --- | --- |
| **(O1)** Anti-tumor CD8+ T cells are shunted to the infected site. | Tumor-specific CD8+ T cells of infected hosts were significantly reduced on day 6 in the TME compared to uninfected hosts and found at high levels at the site of infection but not observed in tissues unrelated to the tumor challenge or infection. | **(O1-M1)** Low-affinity immunological synapses formed between TCRs on anti-tumor CD8+ T cells and self-antigens on tumor cells lead to the lack of arrest of the anti-tumor CD8+ T cells in the TME. |
|  |  | **(O1-M2)** Infection-induced chemokines and cytokines amplify the tumor's ability to egress anti-tumor T<sub>EFF</sub> from the TME. |
|  |  | **(O1-M3)** Non-specific cardiovascular edema caused by infection-induced inflammation affects anti-tumor T<sub>EFF</sub> trafficking. |
|  |  | **(O1-M4)** Infection-induced chemokines chemoattract anti-tumor T<sub>EFF</sub> to the infected lung. |
|  |  | **(O1-M5)** Infection-induced IL-2 retains all types of T<sub>EFF</sub> in the infected lung. |
|  |  | **(O1-M6)** Infection-induced cytokines amplify expression of endothelial PD-L1, which in turn leads to paralysis of anti-tumor T<sub>EFF</sub> in the inflamed lung due to high-affinity PD-1:PD-L1 bonds. |
| **(O2)** Cancer does not suppress the immune system anti-viral response which is capable at the same time of eradicating concomitant infections efficiently. | Cancer does not alter significantly the natural clearance of infection. Influenza infection also does not alter the natural clearance of VACV or the proportion of VACV-tetramer+ CD8+ T cells at the site of influenza infection. | **(O2-M1)** High-affinity immunological synapses formed between TCRs on anti-infection CD8+ T cells and nonself-antigens on infected cells lead to the arrest of the anti-infection CD8+ T cells inside the infected sites until a full clearance of the infection antigen. |
| **(O3)** Therapeutic PD-1 blockade reverses infection-mediated anti-tumor response disruption. | PD-1 blockade decelerates tumor growth in influenza-infected mice as well as rescues the percentage of anti-tumor CD8+ T cells within the TME. | **(O3-M1)** αPD-1 blockade shifts the dynamic equilibrium of the dynamically formed PD-1:PD-L1 bonds towards unbound forms of both PD-L1 and PD-1, allowing anti-tumor T<sub>EFF</sub> to recover from immunologic paralysis and to gain motility. |
|  |  | **(O3-M2)** Due to PD-1-induced mobility reactivated anti-tumor CD8+ T cells may recirculate back to the TME with constitutive lymph motion. |
| | | **(O3-M3)** αPD-1 blockade reactivates PD-1-blocked signaling pathways leading to (i) improved killing capability, (ii) proliferation, (iii) suppression of PD-1 expression, (iv) protection against exhaustion, *etc.* in reactivated CD8+ T cells. |

<sup>a)</sup> Literature citations are directly inserted through the main text.

**(O1) Anti-tumor CD8+ T cells are shunted to the lung during influenza infection.** Here, we discuss six mechanisms (O1-M1) - (O1-M6) from Table 1. In order to get insight into mechanism (O1-M1), we subdivide mechanism (O1-M1-B) into two complementary immunological mechanisms (O1-M1-A) and (O1-M1-B).

#### Mechanism (O1-M1-A)

Cytotoxic CD8+ T cells (T_EFF_) are highly dynamic once activated. They have been shown to rapidly traffic in tissue in response to chemokines and cytokines (Kindt et al., 2007), including CXCL9, CXCL10, INF*γ*, *etc.* (Ogawa et al., 2002; Baaten et al., 2013; Oelkrug and Ramage, 2014; Kim and Chen, 2016; Spranger, 2016; Stein et al., 2016) with a primary function to find and kill target cells expressing cognate antigen (Ag) (Bhat et al., 2014). Dynamic speed and travel patterns of T_EFF_ are predominantly influenced by the tissue environment rather than by mechanisms intrinsic to the T_EFF_. Specifically, activated T cells have been shown *in vivo* to traffic to inflamed skin, even in the absence of cognate Ag (Biotec and Gladbach, 2011), suggesting that Ag alone does not play an essential role in the recruitment of circulating CD8+ T cells (Van Braeckel-Budimir and Harty, 2017).

To this end, a natural question arises: “Why is it the anti-tumor T_EFF_ cells that are shunted to the infected lung, and not vice versa, that is, why is it not the anti-influenza T_EFF_ cells that are shunted to the tumor compartment instead?”

#### Mechanism (O1-M1-B)

The above question can be addressed by reviewing *in vivo* studies describing T cell *arrest* in tissues in direct proportion to the amount of Ag present (Beattie et al., 2010; Deguine et al., 2010; Celli et al., 2011). Indeed, the time needed for the T_EFF_ killing of highly antigenic cells *in vivo* through the cell-to-cell attachment and TCR-pMHC (Ag) binding events can be partiality attributed to effective half-life or “confinement time” of a TCR-pMHC interaction (Aleksic et al., 2010). The process can last for long periods of time depending on the context, (*i*) in the range of 40 minutes, (*ii*) over 3-6 hours and (*iii*) up to 48 hours, in order to form “stable immunologic synapses” (Grakoui et al., 1999; Liechtenstein et al., 2012; Xie et al., 2013; Ortega-Carrion and Vicente-Manzanares, 2016), which are needed to complete a series of signaling events, including co-receptor recruitment and TCR phosphorylation (McKeithan, 1995; Garcia et al., 2007; Breart et al., 2008; Tkach and Altan-Bonnet, 2013; Liu et al., 2014; Das et al., 2015; Parello and Huseby, 2015; Rendall and Sontag, 2017).

Here, the definitions of “long periods of time” and “stable immunologic synapses” should be understood dynamically and not statically in terms of “fast association and dissociation rates” (Tkach and Altan-Bonnet, 2013) that define the “confinement time” of a TCR-pMHC interaction (Aleksic et al., 2010). Multiple studies have indicated that T cells integrate these discontinuous antigen contacts over time and respond in proportion to the cumulative duration of TCR signaling as reviewed in (Tkach and Altan-Bonnet, 2013).

Many tumor-specific antigens provoke only weak immune responses, which are incapable of eliminating all tumor cells (Aleksic et al., 2010). This is in line with the McKeithan-Altan-Bonnet-Germain kinetic proofreading model (KPL-IFFL) (Hopfield, 1974; McKeithan, 1995; Altan-Bonnet and Germain, 2005; François et al., 2013; Courtney et al., 2017), which is based on the well-established fact that T cell activation is selected by evolution to discriminate a few foreign peptides rapidly from a vast excess of self-peptides (François et al., 2013). We use the abbreviation KPL-IFFL for the kinetic proofreading model coupled with limited signaling and incoherent feedforward loop (Fig. SI-2.1) as suggested by (Lever et al., 2016).

Because many tumor antigens are self antigens, often called Tumour-Associated Antigens (TAAs), (Liechtenstein et al., 2012), anti-tumor TCRs may be of lower affinity due to their selection and training against “self”-antigen reactivity (Hogquist and Jameson, 2014) compared with those TCRs evolved to recognize viral epitopes (Irving et al., 2012; Vonderheide and June, 2014). Indeed, the affinity of TCR clones for novel or not previously encountered antigens, like tumor antigens, is remarkably low, typically 1 - 10 *µ*M (Courtney et al., 2017).

This phenomenon is known as “antigen discrimination” (Galvez et al., 2016; Rendall and Sontag, 2017), and is currently discussed in terms of the “antigen-receptor (catch bonds) lifetime dogma” (Feinerman et al., 2008; François and Altan-Bonnet, 2016).

In addition to the TCR antigen discrimination, tumors do not express large amounts of cognate Ag on MHCs to keep systemic tolerance and prevent the development of autoimmune diseases (Liechtenstein et al., 2012), compared with viral infection.

Antigen expression by tumor cells thus determines T_EFF_ motility within the tumor (Boissonnas et al., 2007). Such mobile T_EFF_ cells can follow collagen fibers or blood vessels, or migrate along blood vessels preferentially adopting an elongated morphology (Boissonnas et al., 2007), when nothing can prevent them from freely moving along high infection-induced chemokine gradients toward the infected and inflamed lung.

We finalize the description of this mechanism by pointing out a very important and relevant study (Poleszczuk et al., 2016) which documents intensive motility of anti-timor T_EFF_ cells when the anti-timor T_EFF_ cells enter and leave the TME back to the bloodstream and lymph multiple times before they get finally arrested and absorbed by the multiple-time revisited TME.

#### Mechanism (O1-M2)

Tumors themselves may induce emigration of tumor-specific CD8+ T cells from the TME by employing chemokines like SDF-1/CXCL12 (Joyce and Fearon, 2015; Kim and Chen, 2016). When the level of CXCL12 becomes greater than the levels of other chemoattractors, CXCL12 acts as chemorepeller (Vianello et al., 2005), that is, at low levels, CXCL12 is a chemoattractor, while at high levels CXCL12 is a chemorepeller. Expression of CXCL12 and its receptor CXCR4 is induced by IFN*γ* (Ogawa et al., 2002; de Oliveira et al., 2013).

Sometimes, this effect is called *fugetaxis* (Vianello et al., 2005). Because IFN*γ* is produced in large quantities by different cell types in inflamed infected sites (Kindt et al., 2007), and then circulates to the tumor site within the bloodstream, it can be concluded that (distant) infection can significantly contribute to the egress of anti-tumor CD8+ T cells from the TME.

#### Mechanism (O1-M3)

Due to non-specific cardiovascular edema effects, caused by infection-induced inflammation (Marchuk, 1997), the general circulation pattern of central memory (T_CM_) and na ï ve T cells (Donnadieu, 2016) throughout the body from blood, across high endothelial venules (HEVs) into lymph nodes, through T cell zones, out via efferent lymphatics, and eventually back into the blood through the thoracic duct is significantly perturbed and is redirected to the site of infection-induced inflammation (Marchuk, 1997).

#### Mechanism (O1-M4)

As mentioned earlier, different cells in infected tissues induce cytokine production (Kindt et al., 2007). Cytokines play multiple roles such as chemoattraction of dendritic cells, macrophages, T cells, NK cells, and promotion of T cell adhesion to endothelial cells (Dufour et al., 2002). To this end, significant levels of both activated influenza-specific and non-specific T cells were found present in infected lung and measured (Toapanta and Ross, 2009).

#### Mechanism (O1-M5)

High levels of IL-2 produced by activated anti-infection CD8+ T cells at the infection site counteract the repelling action of CXCL12 (Beider et al., 2003) in contrast to the opposite effect elicited by CXCL12 in the TME as discussed earlier.

#### Mechanism (O1-M6)

The PD-1 mediated control of immune responses depends on interactions between PD-1 on CD8+ T cells and PD-L1 in tissues, inducing CD8+ T cell motility paralysis via PD-1:PD-L1 stable bonds (Zinselmeyer et al., 2013; Schietinger and Greenberg, 2014). We introduce the “paralysis” mechanism **(O1-M6)** into the context of our studies (Fig. 1) by discussing systematically the following specific questions,

**Figure 1:**
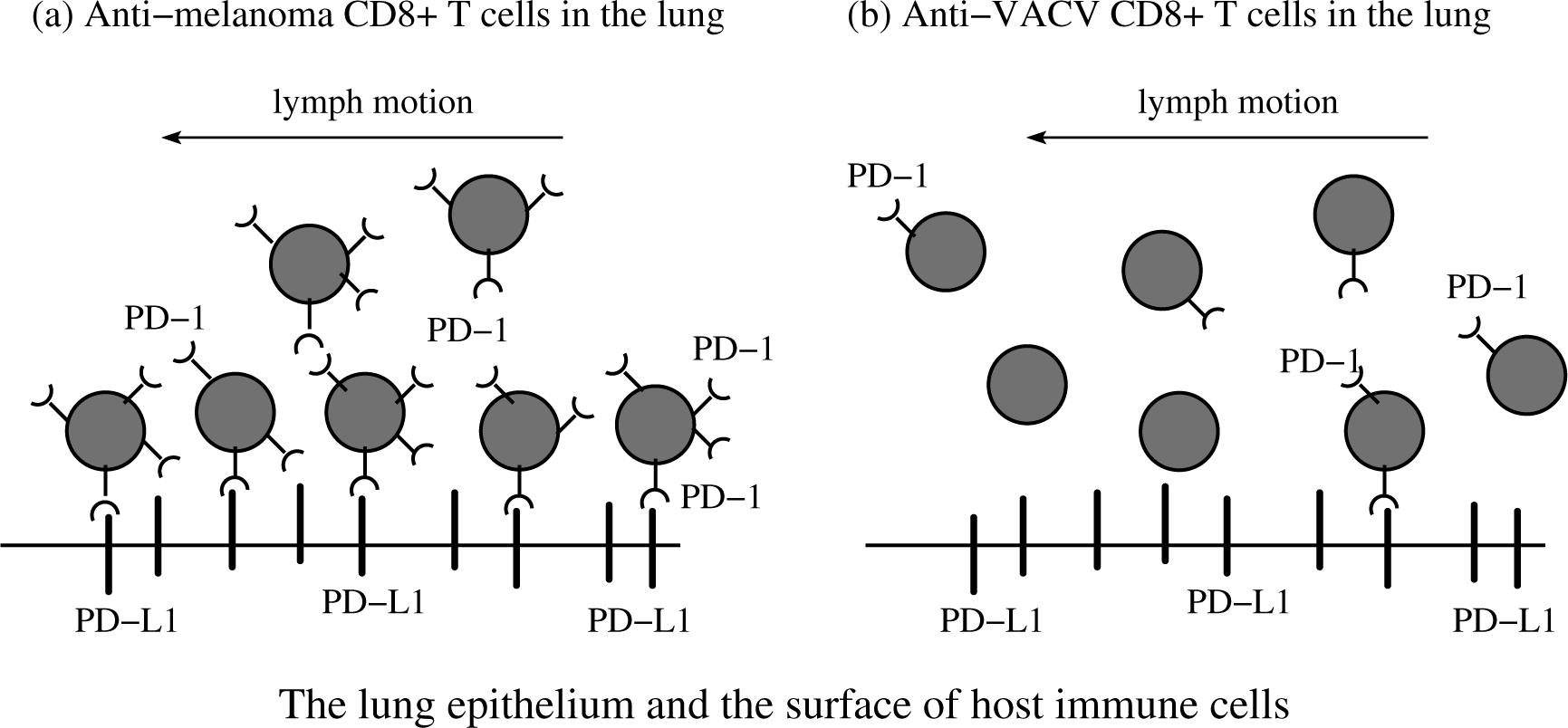
PD-1:PD-L1 induced paralysis of the anti-tumor exhausted CD8+ T cells in the infected site. Panel (a) suggests that anti-melanoma T_EFF_ cells become paralyzed in the infected lung. In contrast, panel (b) suggests that anti-VACV T_EFF_ studied in (Kohlhapp et al., 2016) can freely enter and leave the infected lung with the lymph motion due to the lack of large amounts of PD-1 receptors on their surface. The immune suppressive environment induced by inflammation in the infected lung is caused by multiple interactions between PD-1 receptors, expressed in *large* quantities on the surface of the anti-melanoma T_EFF_, and the PD-L1 ligands expressed in *large* quantities on the surface of various host immune cells (macrophages, DCs, and MDSCs) and the epithelium (Bardhan et al., 2016).

(Q.1) “What triggers expression of PD-1 receptors on CD8+ T cells, and why is the expression triggered in the first place?”

(Q.2) “Why are PD-1 receptors over-expressed in larger quantities on anti-tumor CD8+ T cells and not on anti-influenza CD8+ T cells co-localized within the same infected lung?”

(Q.3) “Why are anti-VACV CD8+ T cells not sequestered in the infected lung when the host is distantly co-infected with both infections, influenza A and VACV infections (in the absence of tumors)?”

To address (Q.1), we follow Simon and Labarriere (2017) who reviewed results highlighting the ambiguous role of PD-1 in defining efficient or inefficient adaptive immune response. Initially, PD-1 transient expression on native T cells is induced immediately upon TCR activation, that is, the number of PD-1 receptors can be regarded as a biomarker of activated and *not* exhausted CD8+ T cells.

The level of PD-1 receptors decreases in the absence of TCR signaling but is maintained upon chronic activation with a persistent epitope target such as in chronic viral infections and in cancer (Wherry et al., 2007; Brown et al., 2010; Pauken and Wherry, 2015b,a). Thus, the number of PD-1 receptors can *also* be regarded as a biomarker of exhausted T cells (Simon and Labarriere, 2017). Importantly, transient expressions of PD-1 and PD-L1 is viewed as a window of opportunity in the combined radiation (RT) and anti-PD-1:PD-L1 therapies (Kosinsky et al., 2018).

The discussed ambiguous role of PD-1, which can be viewed either as a biomarker of activated or exhausted CD8+ T cells depending on the inflammation context, can be explained as follows.

First, although the central immune tolerance mechanism results in the removal of most of the auto- or self-reactive T cells during thymic selection, a fraction of self-reactive lymphocytes escapes to the periphery and poses the threat of autoimmunity. Moreover, it is now understood that the T cell repertoire is in fact broadly self-reactive, even self-centered (Hogquist and Jameson, 2014; Richards et al., 2016).

The strength with which a T cell reacts to self ligands and the environmental context in which this reaction occurs influence almost every aspect of T cell biology, from development to differentiation to effector function (Hogquist and Jameson, 2014). The immune system has evolved various mechanisms to constrain autoreactive T cells and maintain peripheral tolerance, including the constitutive expression of PD-L1 in large quantities in various tissues (*e.g.*, lungs, pancreatic islets, *etc.*), and T cell anergy, deletion, and suppression by regulatory T cells (Sakaguchi et al., 2008; Fife et al., 2009; Francisco et al., 2010; Schietinger and Greenberg, 2014; Bardhan et al., 2016).

Second, although T cells endow their host with a defense that favors pathogen clearance, this efficiency sometimes gives rise to intolerable immunopathology, especially when a pathogen transitions into a state of persistence. For this reason, the immune system is equipped with dampening mechanisms that induce T cell exhaustion via PD-1 and PD-L1 immune regulators (Zinselmeyer et al., 2013; Pauken and Wherry, 2015b; Bardhan et al., 2016). This means that the activated T cells must be attenuated irrespective of whether invaders are eliminated or persist. This is because, quite often, persisting microorganisms may cause less tissue damage than the associated immunopathology as a result of continued lymphocyte cytotoxicity (Speiser et al., 2016). Overall, this means that the immune system prefers to put infection that cannot be eradicated rapidly into a chronic state which should produce less damage to the body than the extended exposition of the body to aggressive CD8+ T cell response.

To address both (Q.2) and (Q.3), knowing more specific molecular and biochemical details are needed, and, so, we postpone the corresponding detailed discussion until Sect. 3.4 and Sect. 4.2, respectively.

##### (O2) Disruption of anti-tumor responses is not due to tumor-induced immune suppression of viral clearance or the inability of the immune system to respond to concomitant challenges

The observation (Kohlhapp et al., 2016) that cancer does not significantly suppress the natural anti-viral response can be explained by similar arguments used to introduce the mechanisms **(O1-M1)** - **(O1-M6)**. These suggest that much weaker inflammation in the tumor site compared with much stronger inflammation in the infected lung may not be enough to force the anti-influenza CD8+ T cells (arrested in the lung as discussed earlier) to egress the lung. The observation (Kohlhapp et al., 2016) that influenza infection did not alter the natural clearance of the VACV or the proportion of VACV-tetramer+ CD8+ T cells at the site of influenza infection can also be explained by the local anti-VACV CD8+ T cells arrest required to kill VACV-infected cells as discussed earlier.

##### (O3) Therapeutic blockade of PD-1 results in reversal of infection-mediated anti-tumor response disruption

Recall that PD-L1 promotes motility paralysis (Zinselmeyer et al., 2013; Schietinger and Greenberg, 2014). In other words, the bond PD-1:PD-L1 mediates locking T cells into a state of prolonged motility arrest by localizing to the environment with abundant PD-L1 expression on stromal cells as discussed earlier, termed “T cell motility paralysis” in (Zinselmeyer et al., 2013; Schietinger and Greenberg, 2014).

Because the bond PD-1:PD-L1 is formed dynamically due to interchanging binding and unbinding processes, blockade of PD-1 shifts the dynamic equilibrium towards dissociation of PD-1:PD-L1 bond, leading to the rapid recovering (about 30 min.) of T cell motility, signaling, and cytokine production (Zinselmeyer et al., 2013; Oelkrug and Ramage, 2014; Pauken et al., 2016). The corresponding details are summarized in Table SI-1.1 of Sect. SI-1.

Reactivated anti-tumor CD8+ T cells then detach from local PD-L1 anchors and start moving with lymph outside of the infected lung and may ultimately return back to the tumor site with the blood flow (Fig. 1), following similar trafficking routes and mechanisms as discussed in (Poleszczuk et al., 2016).

### 3.4 Immunobiochemical reaction network reconstruction

In the previous sections, we developed a functional immunologic reconstruction describing important observations reported in (Kohlhapp et al., 2016). Next, we sought to develop an appropriate minimal reaction network to get mechanistic insight into the corresponding molecular detail of infection-tumor interactions.

#### 3.4.1 Untangling IFFL phenotypic motifs

PD-1 expression on CD8+ T cells is known to be regulated at the level of transcription of its gene *pdcd1* (Lu et al., 2014). More precisely, two upstream conserved regulatory CR-B and CR-C regions are key for PD-1 expression in response to CD8+ T cell activation (Lu et al., 2014). Specifically, TCR signaling induces PD-1 gene expression through the transcriptional activator, Nuclear Factor of Activated T cells, cytoplasmic 1 (NFATc1) (Fig. 2), which binds to CR-C after translocation to the nucleus (Collins and Henderson, 2014; Lu et al., 2014).

**Figure 2:**
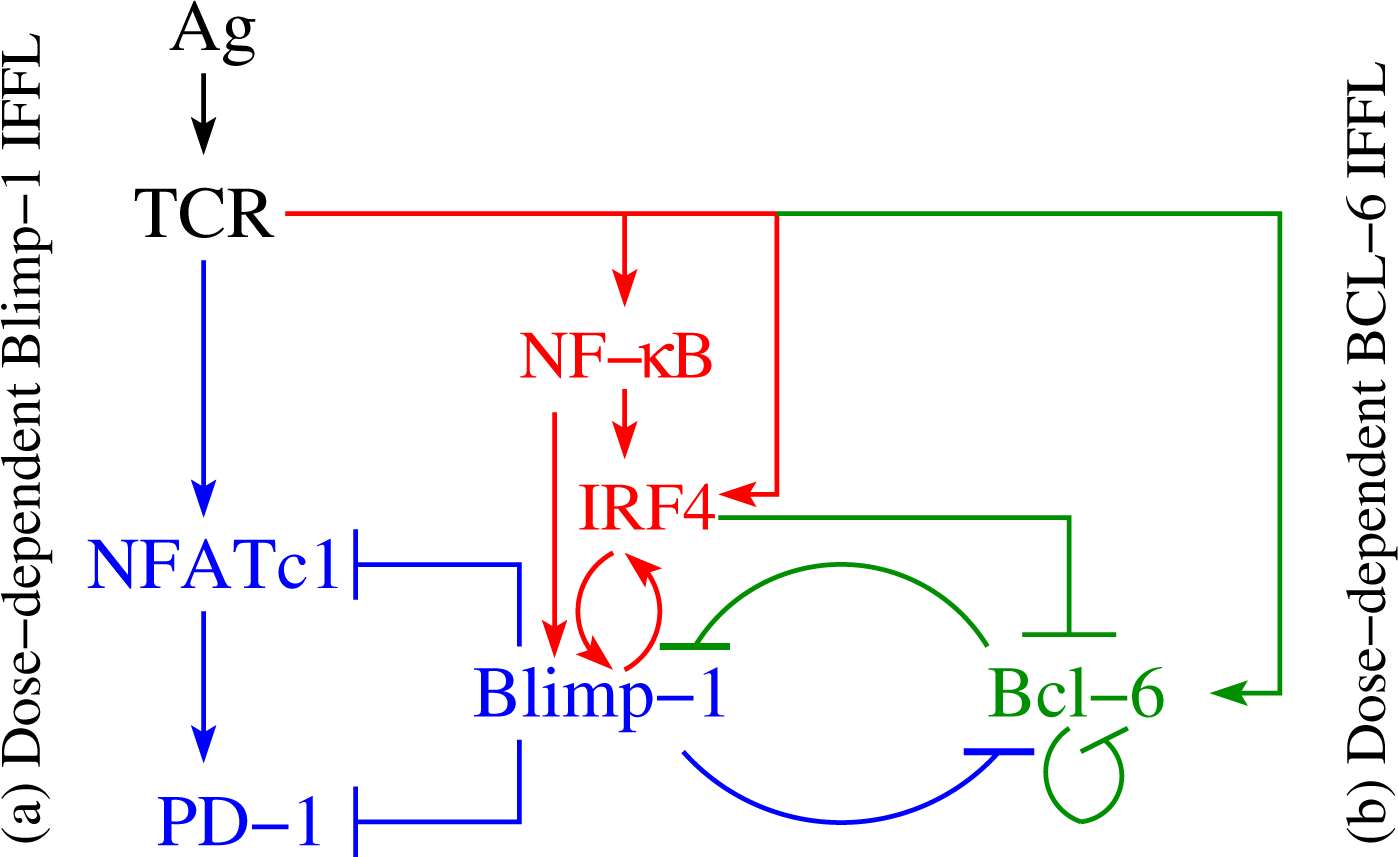
Regulation of PD-1 expression. Two different IFFLs, having a common set of species and regulatory activities highlighted in red, are presented. Both IFFLs are activated by the same input (Ag). Panel (a) depicts a dose-dependent biphasic activation of PD-1. The elements of the corresponding IFFL are highlighted in blue and red colors. When the input, the Ag dose, increases, the output, the PD-1 level, first also increases but then subsequently decreases. Panel (b) corresponds to a dose-dependent activation of Bcl-6. The elements of the corresponding IFFL are highlighted in green and red colors. Over a certain range of input dose, the Ag level, the output, in this case Bcl-6 level, increases but with a subsequent increase in the Ag dose, the Bcl-6 level eventually decreases.

Next, the down-regulation of PD-1 during acute infection (Wherry et al., 2007) suggests that there exists a mechanism that directly represses its expression after initial activation events. Indeed, Blimp-1 (B Lymphocyte-Induced Maturation Protein 1) (Martins and Calame, 2008) has been found to be induced during the later stages of CD8+ T cell activation and was shown to be required for the efficient terminal differentiation of effector CD8+ T cells (Lu et al., 2014). When Blimp-1 is suppressed, the same data suggest that in the absence of Blimp-1, PD-1 expression is maintained by NFATc1 (Fig. 2).

For the sake of completeness, we recall that the existing data also suggest that Blimp-1 represses PD-1 gene expression in CD8+ T cells using three distinct molecular mechanisms (Lu et al., 2014):

1. regulation of the expression of PD-1’s activator NFATc1;
2. alteration of the local chromatin structure; and
3. eviction of the activator NFATc1 from its site that controls PD-1 expression.

In addition, Blimp-1 has been found to be a transcriptional antagonist of proto-oncogene Bcl-6 (B cell lymphoma 6 transcription factor), and vice versa (Fig. 2) (*i.e.*, Blimp-1 and Bcl-6 are known to mutually repress one another) (Calame, 2008; Sciammas et al., 2011; Singh et al., 2014; Daniels and Teixeiro, 2015; Speiser et al., 2016). Specifically, Blimp-1 can bind to the *bcl6* promoter (Shaffer et al., 2002).

Although it is not known exactly how Bcl-6 inhibits Blimp-1 in T cells, it is well known that in B cells Bcl-6, in association with a corepressor MTA3, represses *prdm1* by binding to sites in *prdm1* intron 5 and intron 3 (Shaffer et al., 2000; Calame, 2008; Pasqualucci et al., 2011). Bcl-6 binds its own promoter and inhibits its own transcription (Fig. 2), thus implementing an autoregulatory loop (Pasqualucci et al., 2003; Martinez et al., 2012) (Fig. 2).

Competing with Bcl-6 for intron 5 in *prdm1*, IRF4 (Interferon Regulatory Factor) (Sciammas et al., 2006; Shaffer et al., 2009; Man et al., 2013; Iwata et al., 2017) is shown to be a direct activator of *prdm1* (Fig. 2) by binding to a site in intron 5 (Calame, 2008). At the same time, IRF4 directly represses gene *bcl-6* by binding to a site within its promoter (Calame, 2008; Shaffer et al., 2009), which is rich in IRF4-binding sites (Martinez et al., 2012).

Because IRF4 is known to be activated both directly via TCR and by NF-*κ*B (Boddicker et al., 2015; Vasanthakumar et al., 2017), we have then sought to determine who activates NF- *κ*B in this context and found that NF-*κ*B is activated by TCR signaling (Calame, 2008; Ahmed and Nandi, 2011; Paul and Schaefer, 2013; Daniels and Teixeiro, 2015). Several potential NF-*κ*B binding sites in the *prdm1* promoter have been suggested. It is also known that IRF4 can bind to its own promoter, supporting a positive feedback mechanism by which high IRF4 expression can be maintained (Shaffer et al., 2008; Martinez et al., 2012).

There are additional signaling routes leading to the activation of IRF4 (*e.g.*, via Akt-mediated pathways) which are not discussed here (Calame, 2008).

After a careful analysis of the reconstructed molecular interactions, we have come to the conclusion that this intricate reaction network consists of two subnetworks, see the explanations provided in the legend of Fig. 2. Both subnetworks have the same input from the activated TCR, while the outputs of the subnetworks are different. Namely, PD-1 is the output of the subnetwork color-coded in blue and red, while Bcl-6 is the output of the subnetwork color-coded in green and red. The two subnetworks share a number of commons species and interact with one another via repressive interactions mediated by the three key species color-coded in red, (*i*) IRF4, (*ii*) Blimp-1, and (*iii*) Bcl-6.

#### 3.4.2 Application of the TIFFL motif to unfold infection-tumor interactions

To get more functional insight into the reconstructed backbone reaction network (Fig. 2), we “re-discover” our core TIFFL motifs (Fig. 2) built into a more the complex network (Fig. 3) corresponding to all observations reported in (Kohlhapp et al., 2016).

**Figure 3:**
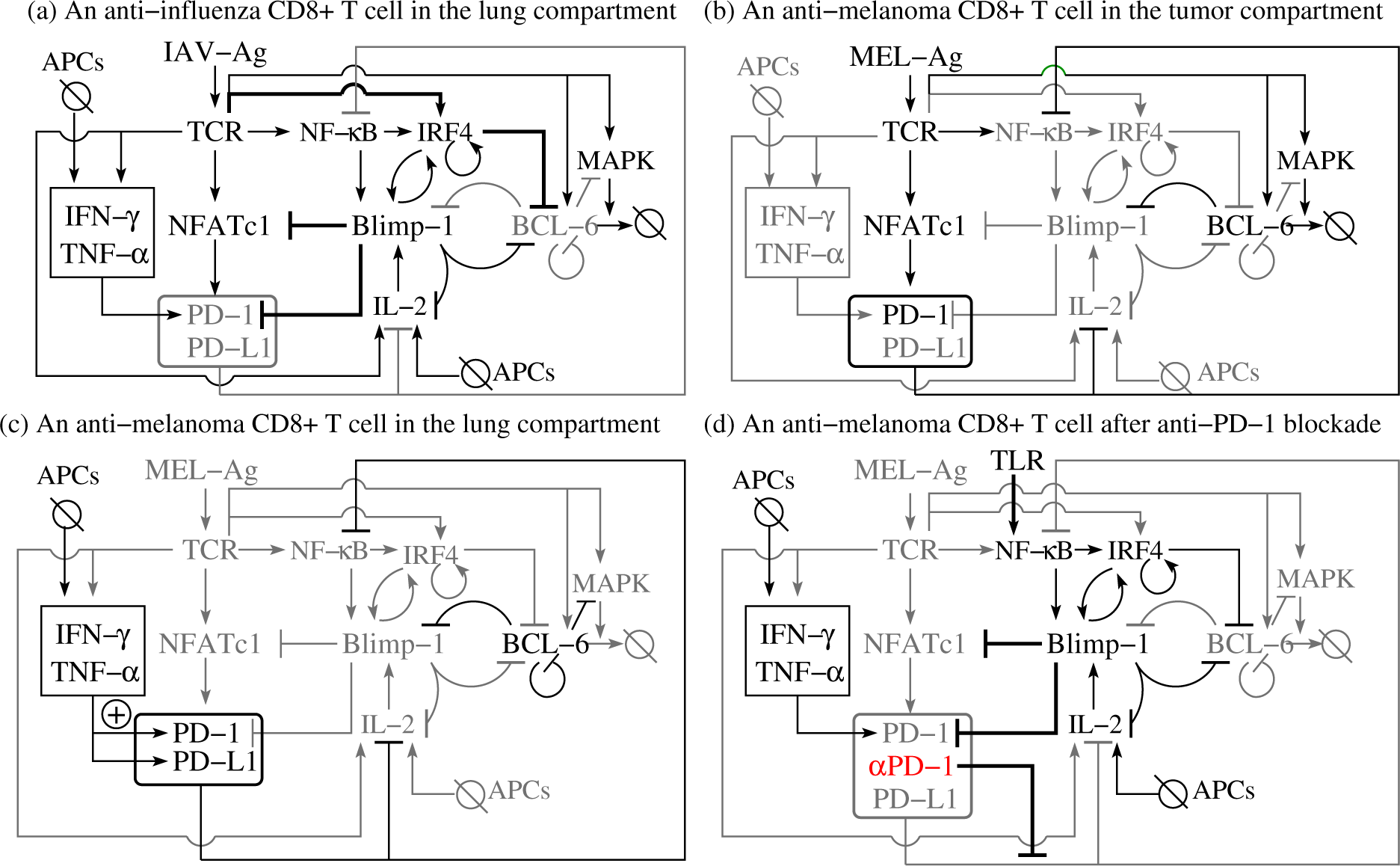
The TIFFL motif in the context of complex influenza-tumor interactions. Panel (a) shows the TIFFL response in an anti-influenza CD8+ T cell in the infected lung. Panel (b) shows the response of the TIFFL circuit in an anti-tumor CD8+ T cell in the TME. Panel (c) shows the TIFFL response in an anti-tumor CD8+ T cell in the influenza-infected lung. Panel (d) shows the TIFFL response in an anti-tumor CD8+ T cell in the influenza-infected lung after PD-1 blockade. Gray color corresponds to weak or disabled reactions shaped by the given inflammation context. Symbol + inside a circle in panel (c) shows the additional PD-1 activation route initiated by external cytokines in the case when the Blimp-1 mediated repression of PD-1 expression is absent. This route does not play any significant role in the case when the expression of PD-1 is suppressed by active Blimp-1 as in panel (a). The abbreviation APCs stands for (influenza) Antigen Presenting Cells.

**Panel (a) of Fig. 3** shows a biochemical reaction network reconstruction customized for the case of an anti-influenza T_EFF_ in the presence of large amounts of cognate Ag in the infected lung. In this case, the immunological complexity of interactions involving cytokines is already overwhelming (Roshani et al., 2014; Dittrich et al., 2015; Duvigneau et al., 2016; Lasfar et al., 2016; Minn and Wherry, 2016; Odenthal et al., 2016). For example, IL-2 activates and is simultaneously repressed by active Blimp-1 both directly and indirectly (Martins et al., 2008; Collins and Henderson, 2014).

The abundance of the cognate viral Ag in the infected lung leads to strong TCR activation which, in turn, results in the simultaneous activation of Blimp-1 and degradation of Bcl-6 (Sect. 3.4.1) followed by suppression of PD-1 transcription with its subsequent degradation. The biochemical detail is in line with and can explains the downregulation of PD-1 in the presence of acute infection (Wherry et al., 2007).

All this may also explain why anti-infection CD8+ T cells are not exhausted during the first phase of the biphasic response of the TIFFL-circuit (Sect. 3.4.1) despite the fact that by-stander and tissue cells express large amounts of PD-L1 caused by large concentrations of pro-inflammatory cytokines such as INF*γ*.

**Panel (b) of Fig. 3** shows the response of the reconstructed circuit in the TME. Specifically, anti-melanoma CD8+ T cells overexpress PD-1 in the presence of large amounts of tumor-specific cytokines such as IL-6, a well-described regulator of Bcl-6 expression (Speiser et al., 2016). Due to relatively low levels of tumor Ag and a weak self-Ag TCR signal (Richards et al., 2016) of anti-tumor CD8+ T cells, the TCR is not activated strongly enough to activate Blimp-1 and, at the same time, the weak activation of the TCR sets the first phase of the biphasic response of the dose-dependent TIFFL motif (Fig. 2).

Indeed, the TIFFL strongly activates Bcl-6 for small and medium TCR strengths, and weakly activates Bcl-6 for high activity levels of TCR. As a result, Bcl-6 is overexpressed, while Blimp-1 is not expressed in the melanoma TME (Speiser et al., 2016), which leads to overexpression of PD-1 on the surface of anti-tumor CD8+ T cells.

**Panel (c) of Fig. 3** shows the TIFFL in an anti-melanoma T_EFF_ relocated into the infected lung. In this case, the conditions discussed just above to introduce panel (b) play the role of a spark plug that activates the transcription of Bcl-6, which represses *prdm1* even after the relocation of the anti-tumor T_EFF_ into the lung. The T_EFF_ can sense INF-*γ* and TNF-*α*, which are abundant in the infected site, produced by both anti-influenza CD8+ T cells and professional antigen presenting cells (APCs), and which strongly stimulate the expression of both PD-1 and PD-L1 (Liu et al., 2017b).

Because the tumor Ag is absent from the infected lung, the TCR is not ligated, and, hence, all routes leading to the activation of both Blimp-1 and Bcl-6 are disabled. We can thus propose that the major route contributing to the PD-1 overexpression here is mediated by INF*γ*- and TNF*α* abundant in the infected lung. The corresponding PD-1 expression activation route is marked by sign + inside a circle in panel (c) of Fig. 3.

**Panel (d) of Fig. 3** shows the TIFFL in an anti-melanoma T_EFF_ cell in the infected lung after administration of PD-1 (*α*PD-1) blockade. Recall that the NF-*κ*B pathway is downregulated in exhausted CD8+ T cells (Speiser et al., 2016). To this end, the PD-1 blockade (marked by symbol *α*PD-1 color-coded in red) in Fig. 3(d), removes the brake from normal T cell signaling pathways (Sect. 3.3, observation **O(3)**, and Sect. SI-1, Table SI-1.1), leading to overexpression of NF-*κ*B (Pauken et al., 2016; Siefker-Radtke and Curti, 2018). Additionally, NF-*κ*B activation is positively regulated through TNFR (TNF Receptor) and TLR (Toll-like Receptor) sensing TNF*α* and viral materials in the infected lung, respectively (Kumar et al., 2004; Liu et al., 2017a; Sun, 2017).

As discussed earlier, NF-*κ*B activates IRF4 (Calame, 2008), and the latter directly represses Bcl-6 (Calame, 2008). In turn, the repression of Bcl-6 removes the brake from the overexpression of Blimp-1, which then leads to reduced numbers of PD-1 receptors on the surface of reactivated anti-melanoma effector cells. This may allow the reactivated T_EFF_ to become mobile and relocate back to the melanoma TME with the lymph flow and blood circulation as discussed previously (mechanism (**O1-M6**) in Sect. 3.3).

### 3.5 Probing immunobiochemical reconstruction modeling

Our modeling goal here is quite simple. Given the discussed specificity of PD-1 expression with respect to different amounts of antigen available in the medium and different values of TCR affinity in terms of the values of the off-rate constant *k*_off_ for the Ag:TCR bond, we focus on the analysis of the dependence of the levels of key species, Bcl-6, IRF4, Blimp-1, and PD-1, on the two parameters, [Ag] and *k*_off_.

#### 3.5.1 Modeling of TCR activation: a spark plug

The importance of TCR activation modeling for our studies is that the ligation of TCR immediately leads to the expression of PD-1 (Sect. 3.2). Based on mechanism **(O1-M6)**, the TCR can be viewed as a “spark plug” for the PD-1 induced processes considered in this work.

To describe the TCR-activated input into our coarse-grained model, which describes the TIFFL depicted in Fig. 2, we have selected the KPL-IFFL model of TCR-activation (Lever et al., 2016). This model is a part of the integrated model of PD-1 expression (Sect. SI-3).

In the KPL-IFFL model, the (scaled) strength of TCR activation depends on two parameters, (*i*) the (scaled) antigen level Ag and (*ii*) the (scaled) off-rate constant *k*_off_ for the Ag-TCR bond (Sect. SI-2). Specifically, the strength of TCR activation is described by the input function *u*(*α*, *κ*) defined in (SI-2.11), where *α* = scaled Ag and *κ* = scaled *k*_off_. The values of the TCR activation function *u*(*α*, *κ*) are plotted in Fig. 4 for a set of parameter values collected in Table SI-2.1.

**Figure 4:**
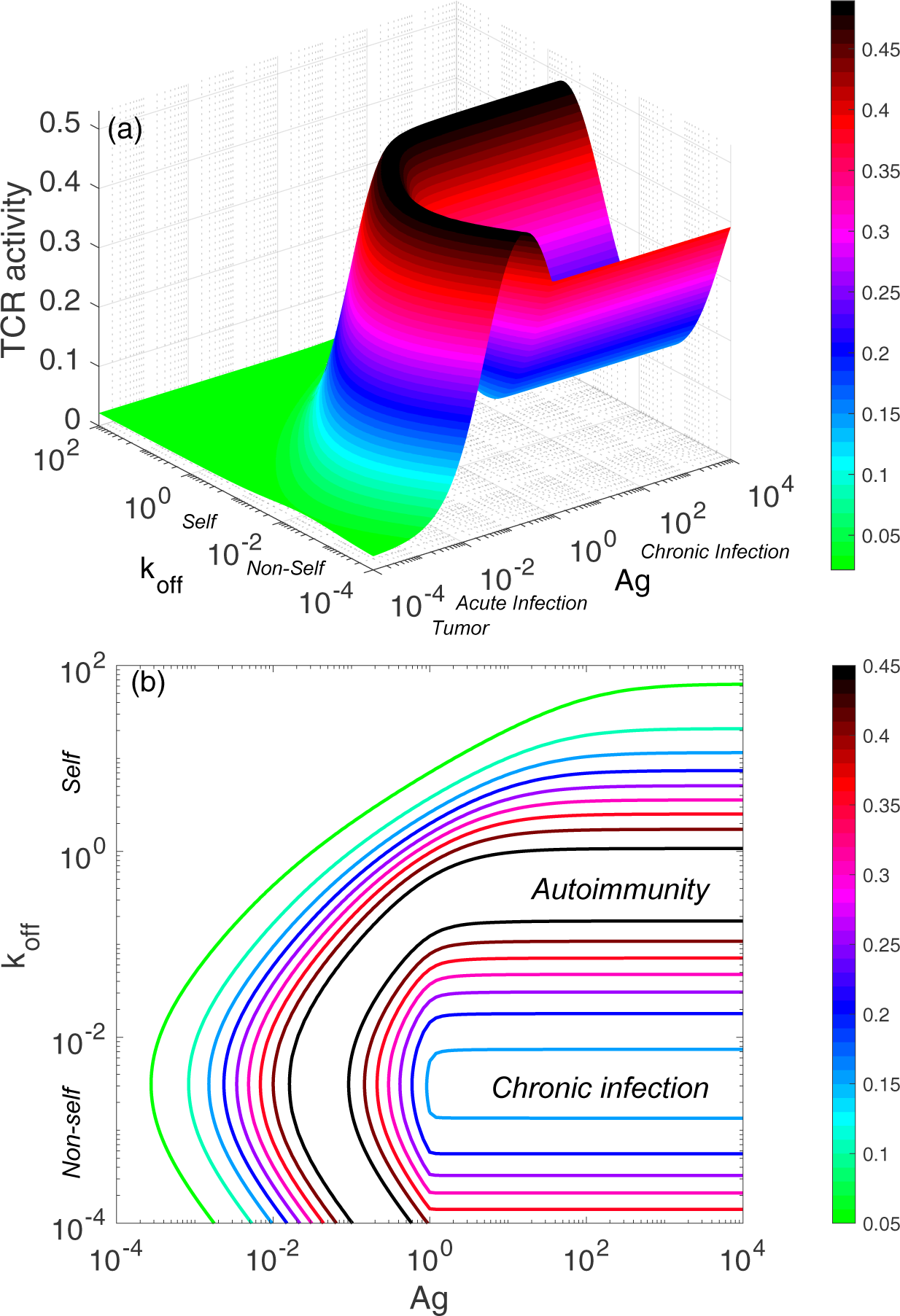
The scaled strength of TCR activation computed from the KPL-IFFL model. Panel (a) shows a 3D-plot of the TCR activity strength depending on the two parameters, the (scaled) concentration of Ag and the (scaled) value of the dissociation constant *k*_off_ for the Ag-TCR bond. Panel shows a 2D-contour plot of the TCR activity strength corresponding to panel (a). Different values of Ag and *k*_off_ are interpreted and classified as corresponding to “*Self*,” “*Non-Self*,” “*Tumor*,” “*Acute infection*,” and “*Chronic infection*” (see the main text for more detail). Colors in the colorbar encode the corresponding values of the function *u*(*α*, *κ*). Both *α* and *κ* are scaled (dimensionless) parameters.

We observe from Figs. 4 and 5 that the TCR activity peaks at optimal values of Ag and *k*_off_, respectively. Such optimal parameter values have been observed in experiments as discussed in (Lever et al., 2016) and rigorously studied in (Rendall and Sontag, 2017).

**Figure 5:**
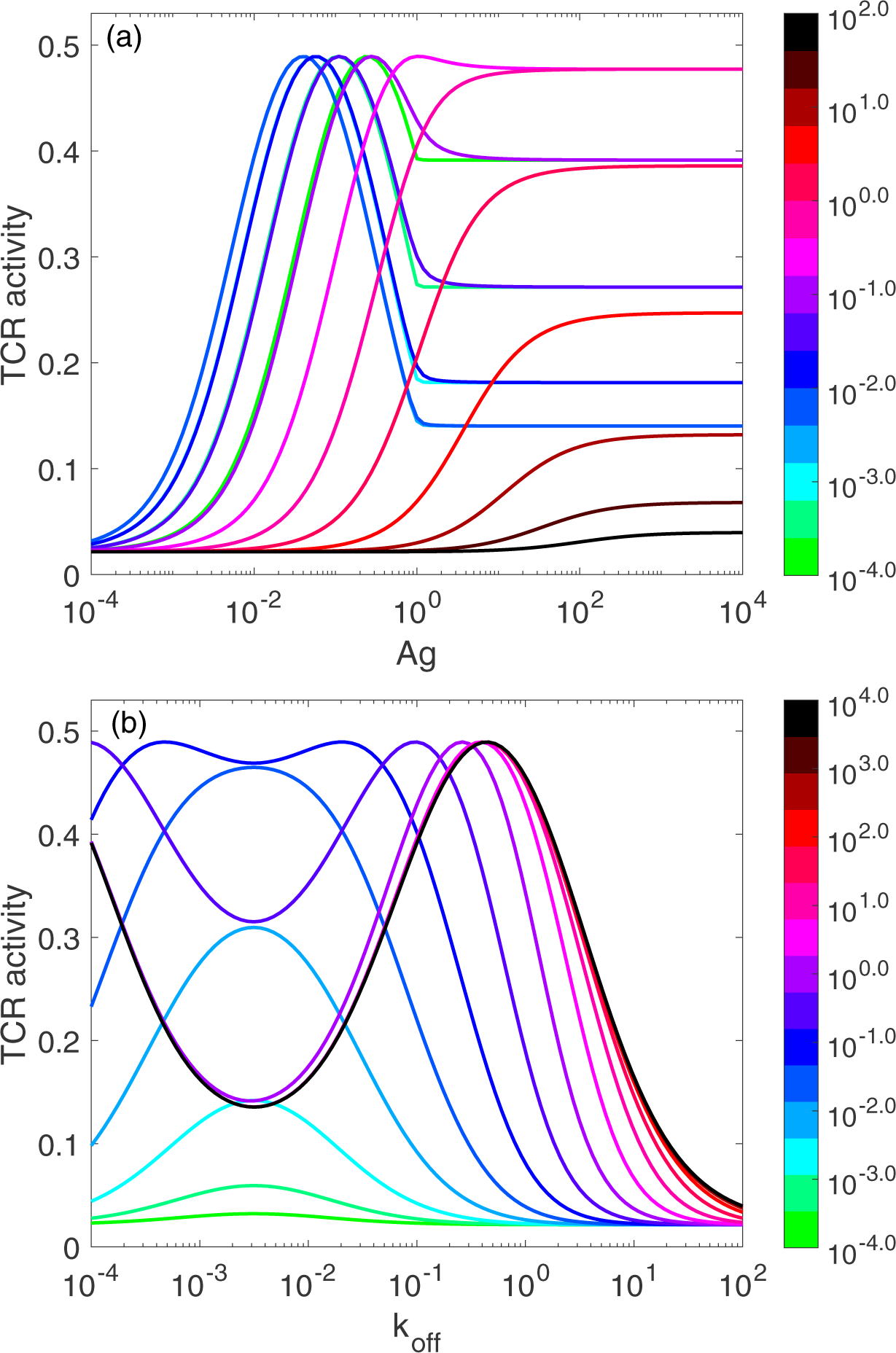
Scaled TCR activty function plots. Panel (a) shows how the TCR activity strength depends on Ag computed for a discrete set of 15 different values of *k*_off_. Panel (b) shows how the TCR activity strength depends on *k*_off_ computed for a discrete set of 15 different values of Ag. In each case, the 15 parameter values have been uniformly selected from the corresponding log_10_ scale, log_10_ *k*_off_ = [-4, 2] in panel (a) and log_10_ [Ag] = [-4, 4] in panel (b). Here and everywhere in the paper, [Ag] is the scaled Ag level normalized by total number of TCRs per a cell, that is, [Ag] = *α*, where *α* is introduced in (SI-2.10f). Similarly, *k*_off_ is a non-dimensionalized parameter, *k*_off_ = *κ*, where *κ* is also introduced in (SI-2.10f).

We believe that the effect of the adaptation (and attenuation) of TCR activation in the presence of very large amounts of Ag corresponding to the TCR activity plateau, coded in blue color in Fig. 4(a) and labeled “Chronic infection” in Fig. 4(b), may be attributed to immune tolerance and adaptive resistance as discussed earlier in mechanism **(O1-M6)** (Sect. 3.3).

The green-blue color coded plots from Fig. 5(a) correspond to this “Chronic infection” region (Fig. 4(b)). These plots can be interpreted as follows. Increasing [Ag] values lead to a sharp increase (in the log-scale) of the TCR activity which peaks at some optimal value of [Ag]. If the antigen (*e.g.*, infection) is eliminated, the TCR activity rapidly returns back to its basal zero level (Fig. 5(a)). However, if the level of Ag continues to increase, the TCR activity reduces to some constant level manifesting immune tolerance to the persistent pathogen as discussed earlier.

We observe from Fig. 4(b) that the optimal (high) values of the TCR activity strength exist within the region (also coded in black color in Fig. 4(a)) for the two parameters Ag and *k*_off_, where both [Ag] and *k*_off_ are *large*. We interpret such regions as corresponding to autoimmunity when certain abnormal combinations of large values of both self-Ag and *k*_off_ may lead to a sharp rise in the activation of self-aggressive T cells. The corresponding plots in Fig. 5(a) are coded using magenta and red colors. In contrast to the “Chronic infection” case, here as the Ag level further increases, the TCR activity does not reduce to a smaller “adapted” level.

The fact that the parameter region marked with label “Autoimmunity” is above the parameter region marked with label “Chronic infection” as seen in Fig. 4(b) can be explained intuitively as corresponding to (*i*) large amounts of Ag in both cases, and (*ii*) different affinities of TCR to antigens: low “self”-affinity corresponding to high values of *k*_off_ and high “non-self” affinity corresponding to low values of *k*_off_.

#### 3.5.2 Modeling of IRF4 rheostat function

In the immunologic literature, IRF4 is called an indispensable molecular “rheostat” that “translates” TCR affinity into the appropriate transcriptional programs that link metabolic function with the clonal selection and effector differentiation of T cells (Man et al., 2013). Because IRF4 plays such a fundamental role, we next model the TIFFL (Fig. 2) at different values of *k*_off_ which defines the TCR affinity (Sect. 3.5.1) to get a more quantitative insight into the IRF4 function.

To simplify the analysis of the multi-parameter model (SI-3.1), we consider here an ideal situation with the PD-1:PD-L1 interaction absent from the model by setting ϕ_*L*_(*P*) = Φ_*L*_(*P*) ≡ 1 corresponding to the condition *L* = 0 in both equations (SI-3.2) and (SI-3.3).

Typical plots for the scaled levels of PD-1, Bcl-6, Blimp-1, and IRF4 in the absence of the PD-1:PD-L1 interaction and at the value of *k*_off_ corresponding to a non-self (foreign) antigen (Sect. 3.5.1) are shown in Fig. 6, see also Figs. SI-4.1 and SI-4.2.

**Figure 6:**
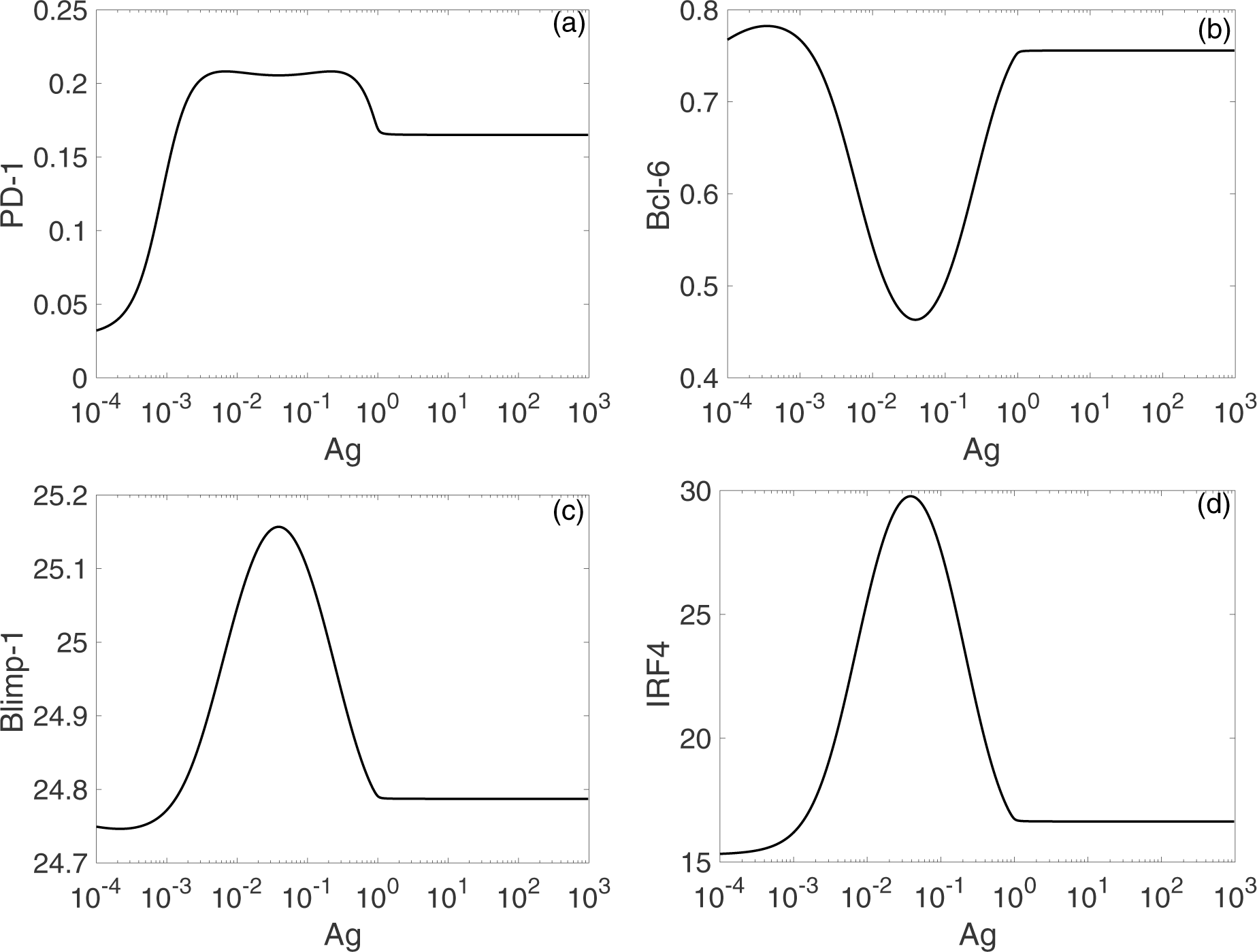
Ag dose-dependent adaptation in the TIFFL. Panels (a) - (d) correspond to species PD-1, Bcl-6, Blimp-1, and IRF4, respectively. The TIFFL is depicted in Fig. 2. All model parameters are selected from Table SI-3.1 with *k*_off_ = 2.43× 10^-3^. To observe an adaptation of the TIFFL with respect to increasing Ag-levels, approximately a 10^3^-fold increase in the Ag-levels is required.

We observe from Fig. 6 that the level of PD-1 (panel (a)) becomes rapidly elevated already at very small values of the scaled concentration [Ag], consistent with experimental observations (Sect. 3.2). A further increase in the scaled concentration [Ag] results in the formation of the PD-1 level plateau, followed by a drop in PD-1.

The increase in the level of PD-1 (Fig. 6(a)) is fully stopped as soon as the level of Blimp-1 (panel (c)) reaches the threshold sufficient to suppress PD-1 expression initiated by TCR activation. We interpret the top (left) plateau in the level of PD-1 (Fig. 6(a)) as the homeostasis maintained by the TIFFL, while the bottom (right) plateau in the level of PD-1 (Fig. 6(a)) can be interpreted as an adaptation to high levels of Ag, a direct consequence of adaptive properties of IFFLs (Alon, 2006; Ma et al., 2009). This adaptation is not “perfect” as compared with the “perfect” adaption levels of Bcl-6, Blimp-1, and IRF4 discussed below in terms of the definition of perfect adaptation and analyses used in (Chiang et al., 2014).

We further observe that in complete agreement with the theory of IFFLs demonstrating biphasic behavior (Alon, 2006; Kim et al., 2008), the levels of Blimp-1 and IRF4 first increase and then decrease, and, at the same time, the level of Bcl-6 first decreases and then increases, while the level of Ag is constantly increased. Remarkably, the levels of all the three species almost perfectly adapt to their respective original states formed initially at very low levels of Ag, as soon as, the level of Ag becomes high enough to establish adaptation. A similar adaptive phenotype is discussed using an example of a generalized enzyme network, conducted in (Chiang et al., 2014).

We believe that all of these modeling results are in line with the immune tolerance discussed earlier in mechanism **(O1-M6)** introduced in Sect. 3.3. However, as soon as the values of *k*_off_ describing the Ag:TCR affinity increase toward “self-antigen” type values (Figs. SI-4.1 and SI-4.2), the behavior of the TIFFL becomes “abnormal,” when all remarkable adaptive properties are completely lost.

To better see the rheostat role of IFR-4 and its impact on the level of PD-1, we then completely disabled IRF4 by setting the value of the parameter *k*_*b*_ to zero, *k*_*b*_ = 0 in the equation (SI-3.1d). This computational experiment can be termed as an “*in silico* IRF4-knockout”. The corresponding plots of PD-1, Blimp-1, and Bcl-6 levels are shown in Fig. SI-4.3. Surprisingly, the shapes of all plots are preserved, and only the magnitudes of the corresponding levels are changed by a factor of 41 *up* for PD-1, by a factor of 5.3 *up* for Bcl-6, and by factor 100 *down* for Blimp-1. We view the abnormal phenotype caused by the loss of IRF4 as a CD8+ T cell adaptation dysfunction.

Motivated by these mathematical conclusions, we proceeded to verify if IRF4 knockout results were previously reported in the literature, and found that *Irf4*-deficient CD4+ T cells display increased expression of PD-1 and other molecular species associated with T cell dysfunction (Wu et al., 2017), which supports our modeling findings.

The T cell dysfunction linked to IRF4 malfunction is also documented in chronic viral infection and tumor models (Wu et al., 2017), which provides an initial clue to the observed fact that anti-influenza T cells have much less PD-1 expressed compared with anti-melanoma T cells in the same infected and inflamed lung, see observation **(O5)** in Sect. 3.1.

#### 3.5.3 Modeling of PD-1 overexpression induced by PD-1:PD-L1 interactions

In this section, we compare the Ag-dose simulation results of the model at *L* = 0.1 (Fig. 6) and *L* = 0.9 (Fig. 7), respectively, for 12 different values of the parameter *k*_off_.

**Figure 7:**
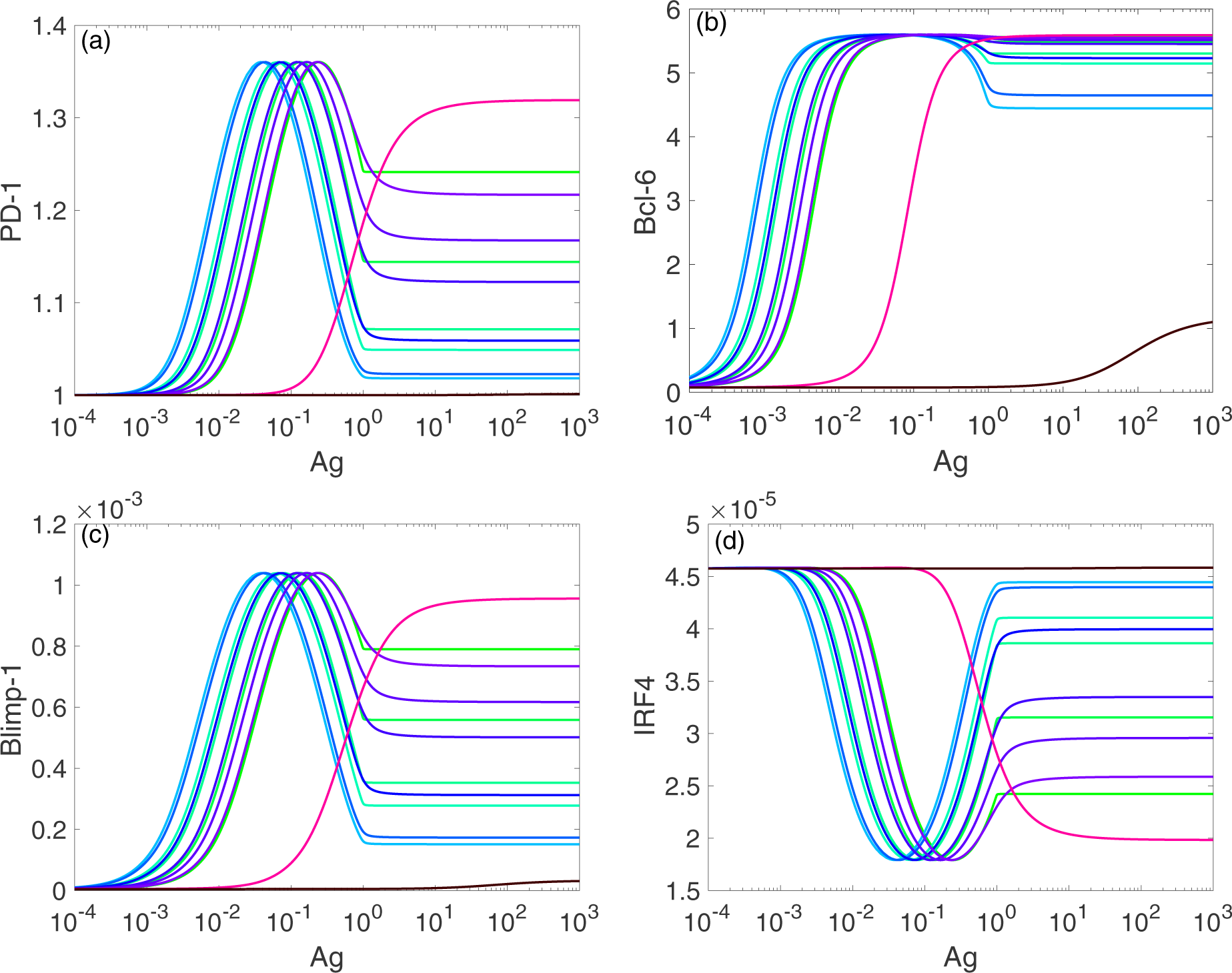
Consequences of the PD-1:PD-L1 interaction and the adaptation effect. The green-blue color-coded plots corresponds to the TIFFL-induced adaptation with respect to increasing Ag-levels. To obtain a full adaptation, approximately a 10^3^-fold increase in the Ag-level is required. The red color-coded plots corresponds to the lack of the adaptation. The panels (a) - (d) correspond to *L* = 0.9. The color-coding schema is as in Fig. SI-4.4. Four different (bottom-up) shades of green color correspond to *k*_off_ = 10^-4^, 2.03 ×10^-4^, 4.13 ×10^-4^, and 5.88 ×10^-4^, respectively. Two shades of blue color correspond to *k*_off_ = 2.43 ×10^-3^ and 7.01 10^-3^, respectively. Four (top-down) shades of purple color correspond to *k*_off_ = 2.03 ×10^-2^, 4.13 ×10^-2^, 5.88 ×10^-2^, and 8.38 ×10^-2^. Magenta color corresponds to *k*_off_ = 1.0. Black color corresponds to *k*_off_ = 49.24.

First, we observe that in the presence of PD-1:PD-L1 interactions, the maximum levels of PD-1 and Bcl-6 are increased by a factor of 6.75 and 7.86, respectively. At the same time, the levels for Blimp-1 and IRF4 are negligibly small, which allows us to interpret that the transcription of these two species is completely suppressed.

From our comparison of the PD-1 level plots in panel (a) of Fig. 6 and Fig. 7, we can conclude that the PD-1:PD-L1 interaction plays the role of an amplifier of transient activation of PD-1 transcription, initiated by the ligation of TCR with Ag presented with an MHC (Sect. 3.2).

Recall that PD-1:PD-L1 interactions terminate the signal transduction pathways, including those pathways that lead to the activation of IRF4 and Blimp-1, by recruiting phosphatases SHP1 and SHP2 (Sect. 3.2). As a result, in the absence of IRF4 and Blimp-1, high levels of PD-1 are sustained even for suboptimal large levels of Ag (Fig. 7(a)).

#### 3.5.4 Modeling of PD-1 expression in anti-melanoma and anti-influenza T cells co-localized in the infected lung

Our last computational experiment compares quantitatively the PD-1 level on the surface of an anti-melanoma CD8+ T cell shunted to the lung with the PD-1 level on the surface of an anti-influenza CD8+ T cell in the lung under the same conditions.

To conduct the computational experiment, the following conditions were taken into consideration: (*i*) the absence of distant tumor Ag in the lung, leading to the shutting down of the TCR signal (*U* = 0), (*ii*) the abundance of inflammatory cytokines, including TNF*α* and IFN*γ*, known to induce the expression of both PD-1 and PD-L1 (Fig. 3(b)), and (*iii*) the abundance of IL-2, which induces Blimp-1 (Fig. 3(b)).

To account for the abundance of the lumped TNF*α*/IFN*γ* species, we have replaced the rate constant *σ*_*p*_ in the equation (SI-3.1b) by the reaction rate expression (SI-3.6).

To account for the abundance of IL-2 in the lung compartment, we have increased the value of the parameter *a*_*b*_ by factor *γ* in the equation (SI-3.1d). In this case, we assumed that IL-2 was secreted by activated T cells (Ahmed and Nandi, 2011) and, hence, IL-2 affected Blimp-1 expression through autocrine and paracrine signaling, depending on the TCR activation strength.

In the case when the value of the parameter *γ* was set to one, the level of PD-1 is increased by a factor of 6 compared with the maximum level of PD-1 shown in Fig. 7 for both anti-influenza and anti-melanoma cases. So, we could conclude that just the TIFFL alone is not enough to counteract the effect of the pro-inflammatory cytokines. Only when a “strong action of IL-2” was taken into consideration by setting *γ >* 5000, the level of PD-1 was suppressed for anti-influenza T cells.

## 4 Discussion

### 4.1 A modeling outlook

In this paper, we have developed a detailed immunobiochemical understanding (Table 1) of complex influenza-tumor interactions (Kohlhapp et al., 2016), formalized in terms of a novel reaction network and its function under four different inflammation contexts (Fig. 3).

We also delineated key detail (Fig. 2) related to other important experimental findings (Kohlhapp et al., 2016), including the expression of PD-1 receptors, in order to form and study a compact mathematical model (SI-3.1) of the phenomenon. Additionally, we employed global sensitivity analysis to quantify and rank the corresponding factors contributing to PD-1 levels, based on the model developed (Sect. SI-5).

Although we did not model spatial effects of shunting anti-melanoma CD8+ T cells from the distant tumor site to the infected lung, and possibly back after PD-1 blockade (Kohlhapp et al., 2016), we did provide a detailed body of experimental evidence available from the literature to explain this phenomenon.

We believe that the modeling of the evolution of cellular populations, including their spatial localization, should be “bi-level,” based on the modeling of trigger-type molecular dynamical variances in each cell. In such a bi-level approach, model (SI-3.1) can be viewed as the “level one” single-cell model before its extension to the “level two” hybrid model of cellular populations observed in (Kohlhapp et al., 2016). In this approach, and in addition to the discussed TIFFL dynamics, one will also need to simulate cellular-level events including: division, death, cell arrest, paralysis, and transition between compartments (Molina-París and Lythe, 2011; Sciammas et al., 2011; Poleszczuk et al., 2016).

Modeling these and similar effects requires a different type of mathematical description based on hybrid approaches, where population and spatial stochastic phenomena could be described by appropriate simple birth and death with immigration formalization, triggered by intracellular molecular interactions (Sciammas et al., 2011), or by more detailed Markov Chain formulations (Poleszczuk et al., 2016). An appropriate stochastic single-cell model can be developed similarly to (SI-3.1) customized to an appropriate Langevin equation with additive noise in parameter values and concentrations. Parameter values in the hybrid population-single cell stochastic model could already be fitted to experimental scatter plots and flow cytometry data provided in (Kohlhapp et al., 2016).

More specifically, the stochastic events discussed can be treated as a first-order process coupled to the fluctuating molecular populations as follows: (*i*) a T cell can be assigned a probability to become exhausted or paralyzed, proportional to its PD-1 receptor concentration; (*ii*) a T cell can be assigned a probability to become arrested to kill infected cells with an antigen concentration and TCR strength greater than the corresponding threshold values; and (*iii*) intervention with *α*PD-1 or *α*PD-L1 blockade should increase signaling leading to reactivation of Blimp-1, followed by putting the brake on PD-1 expression rendering mobility to the paralyzed exhausted T cell again. These probabilities could then be fitted from the data (Kohlhapp et al., 2016) or estimated from the corresponding population fractions if measured accordingly.

### 4.2 Concluding remarks and hypotheses

Motivated by the need to provide a more conceptual and quantitative biology insight into a previously unrecognized immunological factor accelerating tumor growth, an “acute non-oncogenic infection factor” (Kohlhapp et al., 2016), we have analyzed and suggested a number of molecularly detailed immunobiochemical mechanisms to interpret the phenomenon (Table 1).

In our approach, we followed two basic steps: (*i*) first, we suggested an immunobiochemical reconstruction of the TIFFL (Fig. 2) that allowed us to explain experimental data (Kohlhapp et al., 2016) *qualitatively* (Fig. 3), and (*ii*) based on the reconstruction (Fig. 2), we then formed a coarse-grained mathematical model which we used to better understand and interpret the previously unrecognized factor *quantitatively*.

In order to see how “the biological wood emerges form molecular trees” (Gunawardena, 2014) suggested by our TIFFL molecular reconstruction (Figs. 2 and 3), we formulate below a plausible hypothesis to explain the unrecognized factor concisely.

Also, as an unexpected “by-product” of our primary research on the unrecognized factor, we will formulate another hypothesis on a “window of opportunity” previously identified in the context of the combined RT and anti-PD-1:PD-L1 therapies (Kosinsky et al., 2018).

Finally, we will discuss to which extent our mathematical modeling efforts have been required and justified to complete this work, and why just the reconstruction alone was not enough to explain the unrecognized factor phenomenon.

We formulate our first hypothesis on the unrecognized immune factor by suggesting that it is the loss of Ag dose-dependent adaptation of the expression of PD-1 receptors in the anti-tumor CD8+ T cells that is a major factor resulting in multiple effects in the presence of acute non-oncogenic infection, including: (1) shunting anti-tumor CD8+ T cells to the inflamed lung, (2) sequestration of the relocated anti-tumor CD8+ T cells in the lung, and (3) related to (1) and (2) accelerated tumor growth.

Indeed, the TIFFL reconstruction and its model explain a non-monotonic behavior of the levels of PD-1 receptors on the surface of CD8+ T cells as a function of the antigen concentration (Figs. 6 and 7). Specifically, in the case of acute VACV infection, the level of PD-1 receptors on the surface of Ag-experienced anti-VACV CD8+ T cells first increases and then decreases to lower levels in the course of the virus replication (Fig. 8(B)), the hallmark of a fundamental IFFL-induced adaptation effect (Alon, 2006) as also discussed earlier in this work. Therefore, if such cells are trafficking across the infected lung, chances that the cells bearing small numbers of PD-1 receptions will loose their motility due to PD-1:PD-L1 interactions in the infected lung is low (Fig. 3).

**Figure 8:**
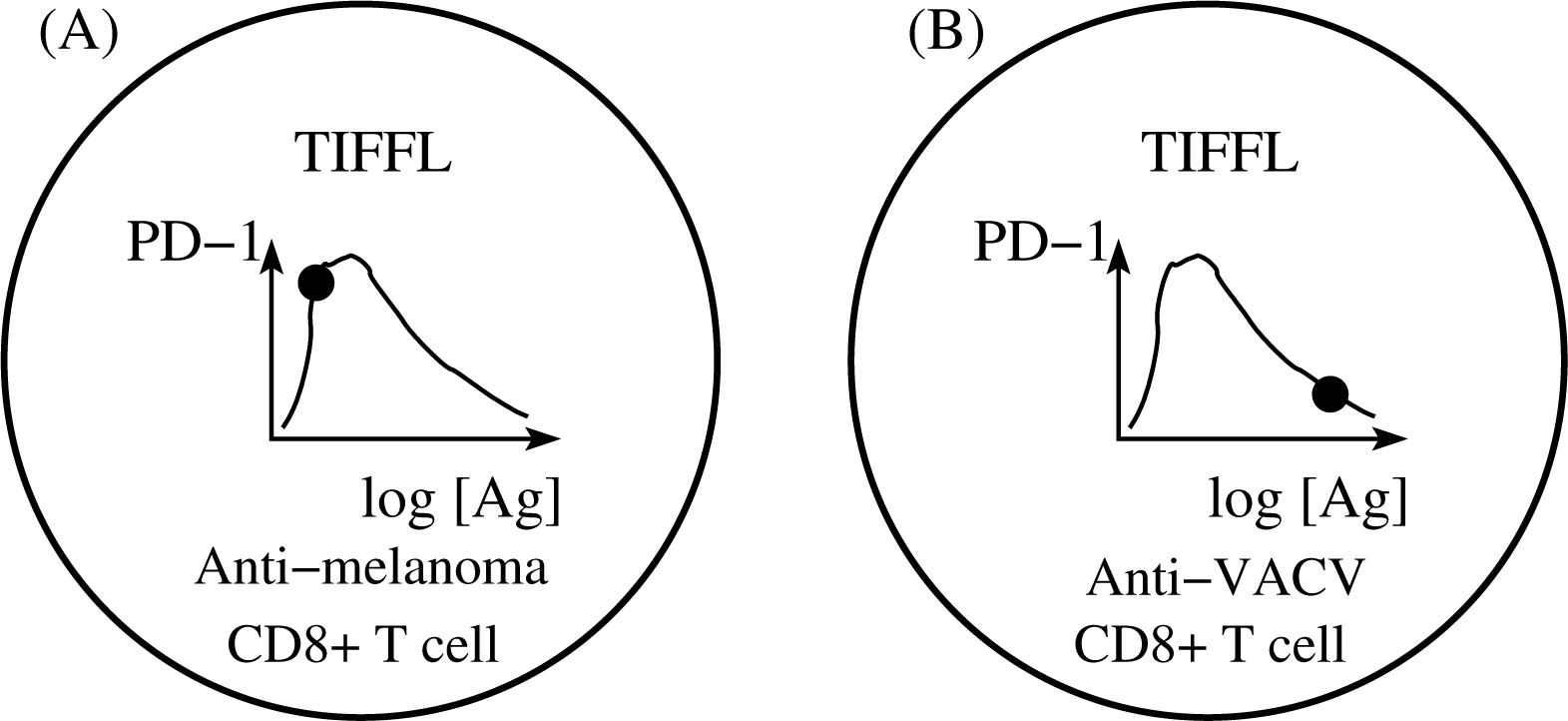
Schematic illustration of Hypothesis 1. Solid filled circles on the corresponding graphs of PD-1 receptor levels, plotted versus the log-concentrations of Ag, correspond to the levels of PD-1 receptors on anti-melanoma (A) and anti-VACV (B) CD8+ T cells, respectively, at the moment when these may enter the influenza A infected lung. State (B) corresponds to a fully developed TIFFL-induced adaptation of the expression of PD-1 receptors with respect to the increasing levels of Ag, while state (A) is characterized by the lack of such adaptation.

In contrast, in the case of Ag-experienced anti-tumor CD8+ T cells, due to both much smaller levels of tumor antigens and lower TCR affinity, the strength of the TCR signal in anti-tumor CD8+ T cells may not be enough to activate Blimp-1 and IRF4 species to suppress PD-1 expression (Figs. 2 and 3). The lack of the expression of Blimp-1 in melanoma is known experimentally (Speiser et al., 2016) as we also discussed earlier in this work. As a result, chances that T cells bearing large numbers of PD-1 receptors (Fig. 8(A)) will be paralyzed in the infected lung due to PD-1:PD-L1 interactions are high. This happens because the amounts of available self-Ag in the TME, and the Ag-TCR affinity are not enough to move the anti-melanoma CD8+ T cell from state (A) to state (B) as shown in Fig. 8.

Importantly, the fact that the higher levels of PD-1 receptors on anti-melanoma CD8+ T cells compared with much lesser levels of PD-1 receptors on anti-influenza CD8+ T cells co-localized in the same infected lung were observed in (Kohlhapp et al., 2016) is totally in line with the hypothesis, in which case the anti-melanoma and anti-influenza CD8+ T cells correspond to states (A) and (B) of Fig. 8, respectively.

As to our second hypothesis on the window of opportunity in the context of the combined RT and anti-PD-1/PD-L1 therapies defined as corresponding to peaked levels of PD-1 receptors on anti-tumor T cells (Kosinsky et al., 2018), we propose that the window of opportunity may mechanistically emerge as function of the TIFFL (Figs. 2). Therefore, we hope that our findings may have broader implications not only in systems immunology of interacting pathogens and tumors, but, additionally, some specific aspects of our molecular detailed immunobiochemical reconstruction can be useful in seemingly unrelated current endeavors in immunotherapy.

Concerning the arguments that justify the use of mathematical modeling in our work, we can say the following. Indeed, as soon as our immunobiochemical reconstruction was completed, and the core circuit (Fig. 2) was recognized as an Incoherent Feed-Forward Loop (Alon, 2006), it became clear that the adaptation of the expression of PD-1 receptors with respect to cognate antigen levels in terms of the corresponding non-monotonic PD-1 levels should play an important role. After we embedded the TIFFL into different inflammation contexts (Fig. 3), the reconstruction helped us better understand the experimental observations *qualitatively*.

However, it was not clear to which extent the qualitative considerations could be helpful in the discrimination of the two cases depicted in Fig. 8 until we developed a mathematical model used to obtain critical computational estimates as follows.

First, as we can easily see from Figs. 6 and 7, already negligibly small amounts of cognate Ag result in a sharp increase in the level of PD-1 receptors on the surface of Ag-experienced CD8+ T cells, which is in a total agreement with the known experimental evidence (Sect. 3.3). At the same time, our quantitative estimates, which can also be obtained from the same Figs. 6 and 7, show that changes in the Ag levels within several order of magnitude (approximately a 10^3^-fold increase counted from the onset of the sharp increase in PD-1 levels) are required for the adaptation to occur, moving the Ag-experienced T cell from state (A) to state (B) as shown in Fig. 8.

While it is reasonable to assume that exponentially replicating viruses can lead to a 10^3^-fold increase in the amount of the cognate Ag sufficient to move Ag-experienced anti-VACV T cells from state (A) to state (B) as in Fig. 8 within 7 - 10 days of the acute infection, it is highly unlikely for tumor cells to divide as fast as the viruses do.

Therefore, it is highly unlikely that during such short periods of time, when acute infection develops and is cleared, the growing and replicated tumor cells can increase the amount of antigen within several orders of magnitude, and, as a result, the majority of anti-tumor T cells will continue to be in state (A) during those short time intervals. An additional important factor here is a weak affinity of TCRs to self-antigens.

Because it is now well understood that Ag-experienced T cells actively circulate across the host’s body with the bloodstream and lymph motion (Poleszczuk et al., 2016), chances that T cells in state (A) will be sequestered in the inflammation regions, while T cells in state (B) will continue to freely circulate through the same inflammation regions are equally high.

Finally, the fact that the coarse-grained model development and simulations require much less time and efforts compared with the time and efforts to be invested into mechanistic reaction network reconstruction further justifies the usefulness of rapid quantitative estimations based on modeling and simulations as done in this work.

## Supporting Information

### SI-1 Reactivation of exhausted effector cells

Because the PD-1 blockade reactivates exhausted anti-tumor CD8+ T cells, sequestered in the infected lung and return, possibly, them back to the TME (Kohlhapp et al., 2016), we briefly summarize relevant known results on the exhausted T cell reactivation (Table SI-1.1). Our summary is based on the recent reports (Zinselmeyer et al., 2013; Pauken et al., 2016; Wang et al., 2017).

**Table SI-1.1:** A brief summary of T_EX_ reactivation after PD-1 blockade.

| Reactivation Effect | Mechanisms |
| --- | --- |
| **(E.1) reactivation of T<sub>EFF</sub> functions in T<sub>EX</sub>:** |  |
| (E.1.1) *improved cell cycle and proliferation* | increased transcription of cell division genes<br>increased levels of Ki-67 |
| (E.1.2) *improved response to antigen* | elevated co-production of INFγ and TNFα |
| (E.1.3) *improved motility and chemotaxis* | upregulated expression of *cxcl9* and *cxcr3* |
| (E.1.4) *improved killing capability* | increased levels of granulocytes |
| (E.1.5) *suppression of PD-1 expression* | upregulated expression of *prdm1* encoding Blimp-1 |
| (E.1.6) *protection against exhaustion* | upregulated expression of *il7r*<br>OCR state specific regulation of *ctla4* |
| **(E.2) reactivation of T<sub>MEM</sub> functions in T<sub>EX</sub>:** |  |
| (E.2.1) *negative regulation of apoptosis* | increased levels of phospho-STAT5 |
| (E.2.2) *improved adhesion* | unknown |
| (E.2.3) *improved regulation of activation* | elevated production of INFγ |
| **(E.3) transient reinvigoration of T<sub>EX</sub> (peaked in 3-weeks)** |  |
| *transient and Ag-dose dependent expression of prdm1 encoding Blimp-1* | small and large amounts of Ag repress *prdm1*<br>medium amounts of Ag activate *prdm1* |
| **(E.4) signaling and immunometabolic effects:** |  |
| (E.4.1) *signaling* | upregulation of genes encoding NF-κB and IRFs |
| (E.4.2) *lipid metabolism* | upregulation of genes encoding PPARγ and RXRα<br>downregulation of *srebp1* |
| (E.4.3) *de-novo cholesterol pathway, and glycolysis* | unknown |

Several gene signatures based on the analyses of populations of dysfunctional CD8+ T cells from cancer and chronic viral infections have been published and reviewed by Wang et al. (2017). These signatures confirm great similarity between virus- and cancer-associated CD8+ T cell dysfunction. Due to these published gene signature comparisons, we believe that Table SI-1.1 further supports mechanisms formulated in the main text.

### SI-2 CD8+ T cell receptor activation input

#### SI-2.1 The KPL-IFFL mechanisms

The input *P* to our model is carried out by the signal initiated by the activated CD8+ T cell receptor (Fig. SI-2.1), which is computed using a mathematical model describing kinetic proof-reading with limited signaling coupled to an incoherent feedforward loop (KPL-IFFL) suggested in (Lever et al., 2016).

**Figure SI-2.1:**
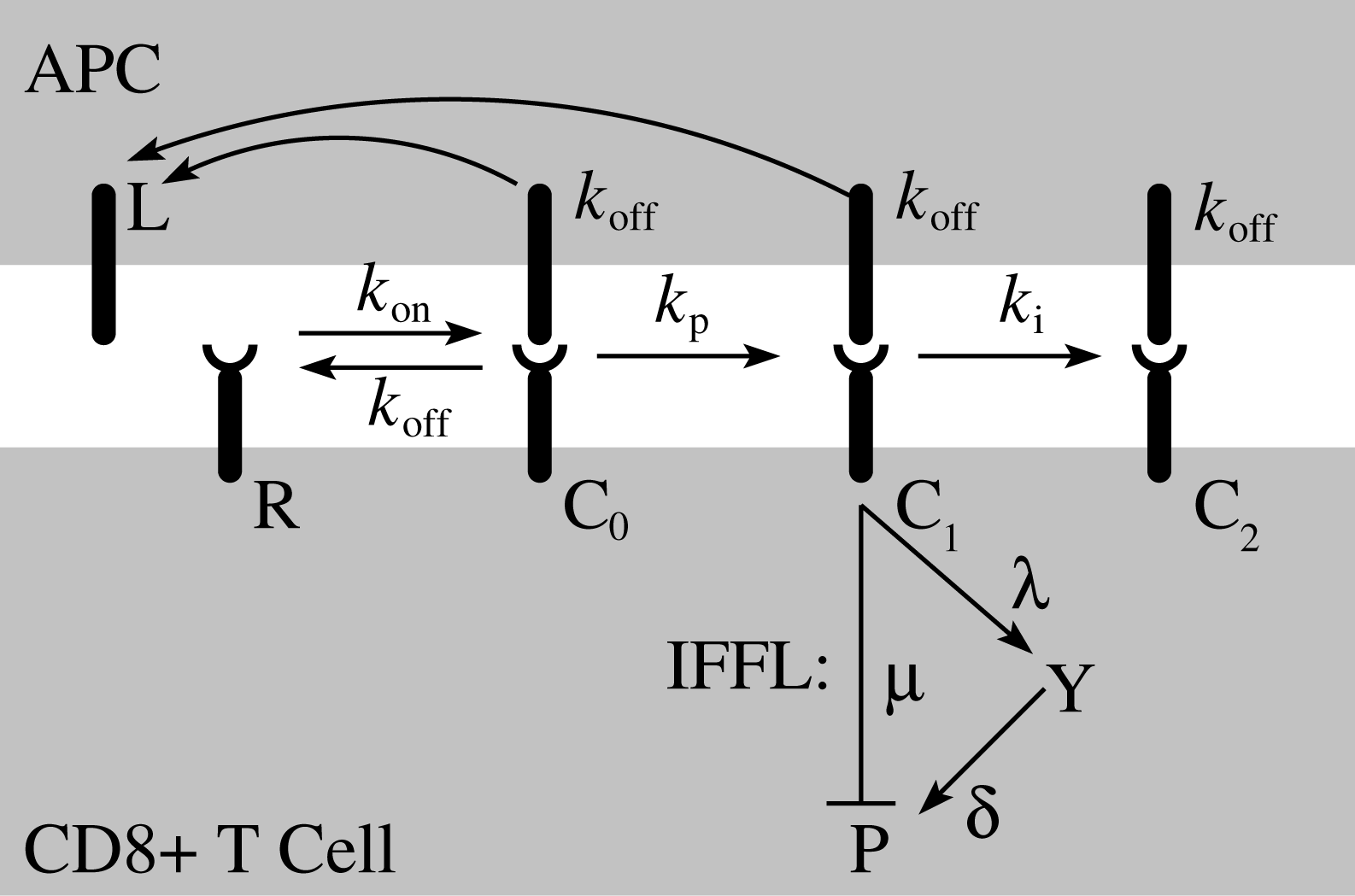
TCR activation via kinetic proofreading with limited signaling coupled with IFFL. After ligation of a T cell receptor (R) with ligand L, presented by a professional antigen presenting cell (APC), a ligand-receptor complex, C_0_, can be formed. The complex C_0_ undergoes a series of kinetic proofreading steps before it reaches the state C_1_ allowed to initiate a downstream signaling. The prolonged duration of the active state C_1_ is penalized by its convertion to the disabled state C_2_. Finally, the active complex receptor gets phoshorylated via a series of reaction steps regulated by positive and negative feedforward and feedback loops, represented by one simplified IFFL with repression (*µ*), amplification (*λ*), and activation (*δ*) parameters. Here, IFFL stands for Incoherent Feed Forward Loop. Explanations of putative proteins Y and P that lump other intermediates and the motivation of the IFFL can be found in (Lever et al., 2016).

#### SI-2.2 The KPL-IFFL model

For the sake of consistency in the integration of the model (Lever et al., 2016) with our model describing the core circuit shown in Fig. 2, we briefly derive functional relationships needed for the models’ integration, adapting the discussion in (Lever et al., 2016).

Specifically, our objective here will be to derive the function *u*(*α, κ*), which we define as a non-dimensionalized input *P* in (SI-2.10b), and for which the final expression is given in (SI-2.11). The scaled function *u*(*α, κ*) depends on two state variables *α* and *κ*, the scaled level of Ag and the scaled value of the off-rate constant *k*_off_, respectively.

A mathematical model (Lever et al., 2016) is

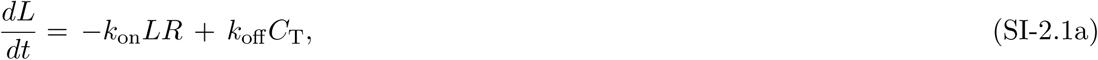

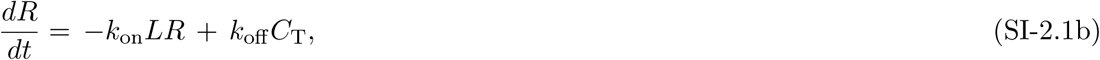

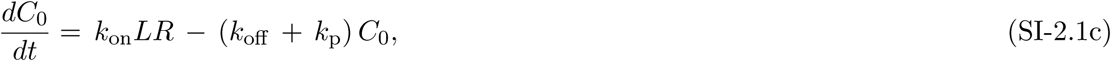

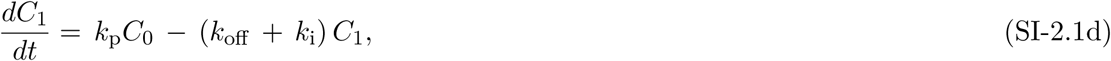

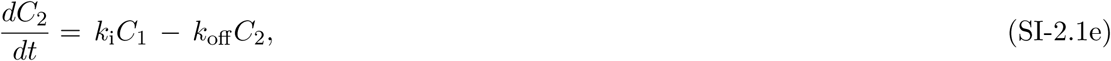

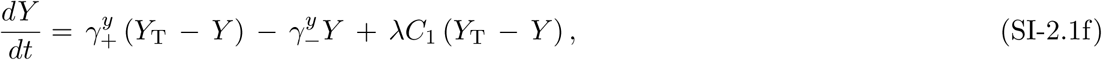

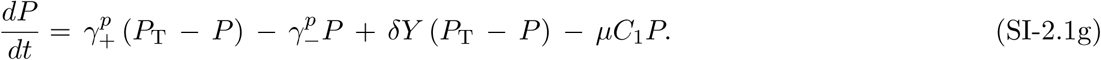

The state variables of the model (SI-2.1) are defined in the legend of Fig. SI-2.1; *k*_on_ and *k*_off_ are on- and off-rate constants, *k*_p_ is the kinetic proofreading rate constant, *k*_i_ is the kinetic rate constant for transforming of the active complex into the inactive complex; *µ*, *λ*, and *δ* are defined in the legend of Fig. SI-2.1; and *C*_T_ is the total number of all ligand-receptor complexes,

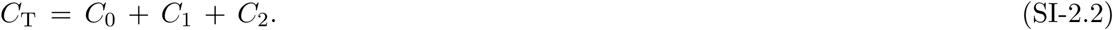

Here, *C*_T_ does not correspond to any conserved moiety and, instead, changes in time.

The putative proteins *Y* and *P* that lump other intermediates (Lever et al., 2016) spontaneously flip between active and inactive states with rate constants 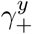 and 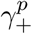, and 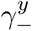 and 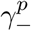, respectively. Additionally, the active protein form *Y* results from the interaction of its inactive form with the activated complex *C*_1_, and the intensity of this process is described by the value of the amplification rate constant *λ*. Similarly, the active protein form *P* additionally results from the interaction of its inactive form with *Y*, and, in turn, the intensity of this process is described by the value of the activation rate constant *δ*.

The reaction network depicted in Fig. SI-2.1 has the following two moiety conservation relationships,

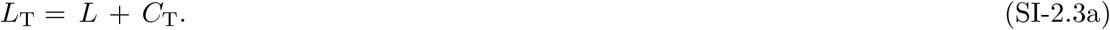

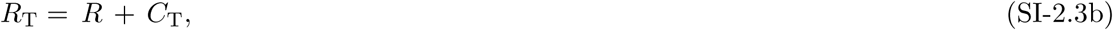

Due to the relationships (SI-2.3b) and (SI-2.3a), the corresponding first two equations (SI-2.1a) and (SI-2.1b) in the model (SI-2.1) become redundant and are omitted from further analysis.

Setting the model linearly independent equations (SI-2.1c) - (SI-2.1g) at steady state, we can obtain the following algebraic relationships,

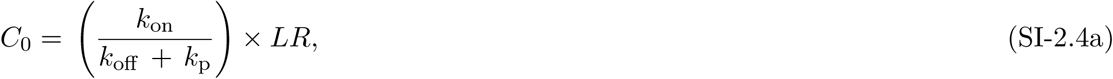

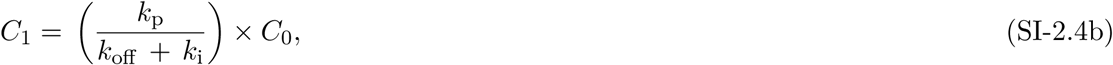

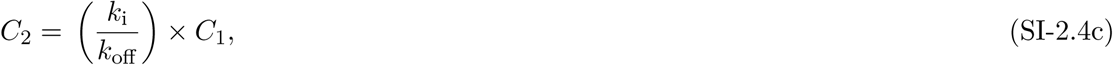

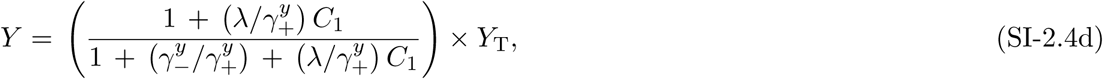

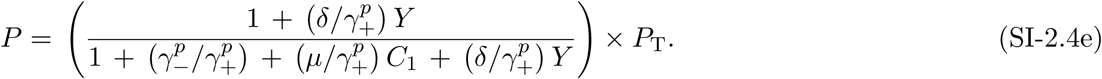

Next, we eliminate the product *LR* from (SI-2.4a) by using (SI-2.4a) - (SI-2.4c) in (SI-2.2),

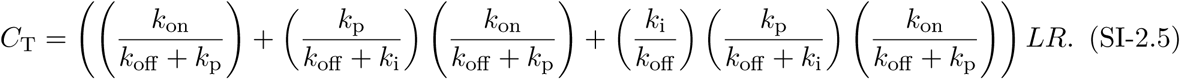

After simple algebraic manipulations, we obtain from (SI-2.5) that

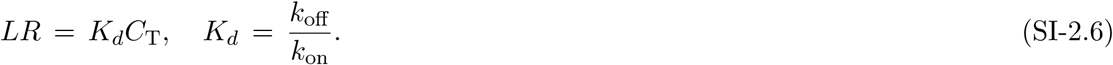

Using (SI-2.6) in (SI-2.4a), and then (SI-2.4a) in (SI-2.4b), followed by using (SI-2.4b) in (SI-2.4c), we obtain

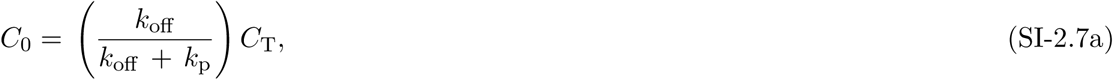

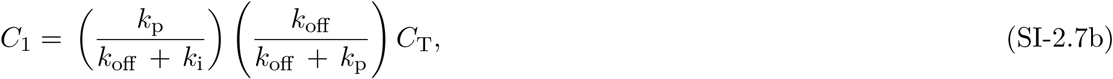

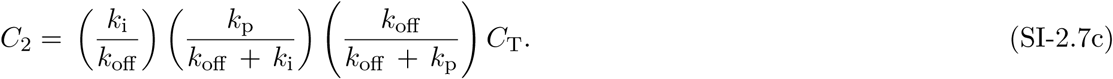

Note that *C*_T_ is still unknown in (SI-2.7). To compute *C*_T_, we use an alternative expression for the product *LR*.

Indeed, we can obtain from (SI-2.3a) and (SI-2.3b) that *L* = *L*_T_ − *C*_T_ and *R* = *R*_T_ − *C*_T_, respectively. Now, using *LR* = (*L*_T_ − *C*_T_) (*R*_T_ − *-C*_T_) in (SI-2.6), we come to a closed quadratic equation with respect to *C*_T_,

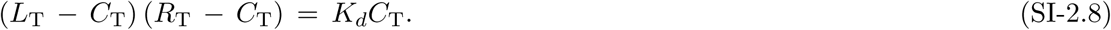

Solving the quadratic equation (SI-2.8) with respect to *C*_T_, we obtain two solutions, only one of which corresponds to the biologically meaningful condition, *C*_T_ = 0 at *L*_T_ = 0,

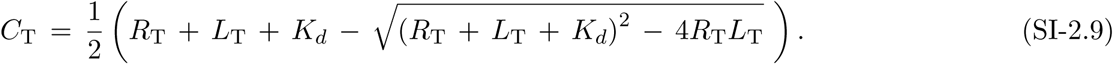

The solution (SI-2.9) also corresponds to the stable equilibrium in the system of linearly independent equations (SI-2.1c) - (SI-2.1g).

It is convenient to nondimensionalize the equilibrium solution of (SI-2.1) given by the expressions (SI-2.4a) - (SI-2.4e), and (SI-2.9) by scaling all state variables and parameters as follows,

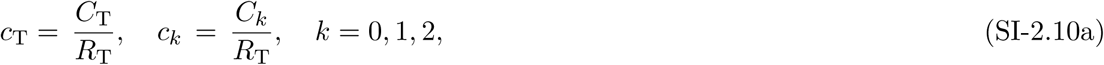

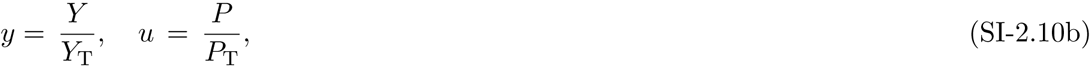

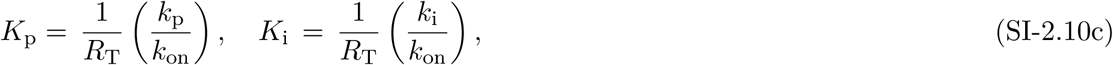

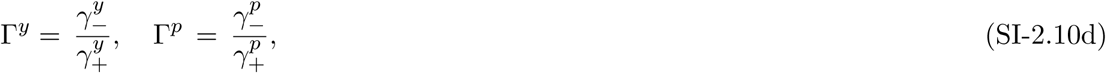

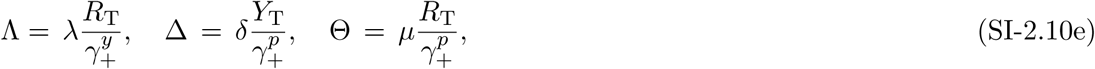

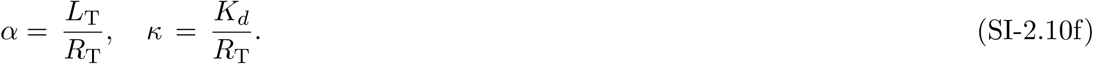

We obtain from the rescaled (SI-2.4e) that

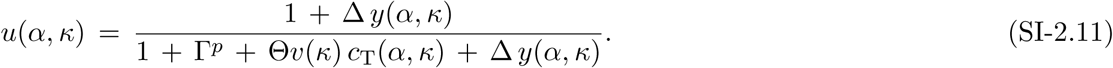

In (SI-2.11), the functions *c*_1_(*α*) and *y*(*α*) are obtained from the corresponding expressions (SI-2.4b) and (SI-2.4d) rescaled as discussed earlier,

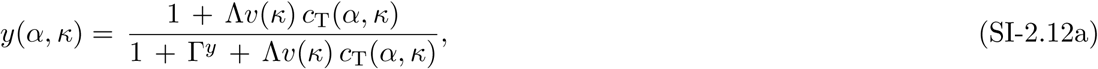

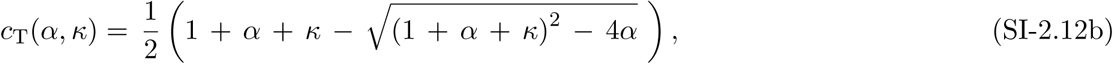

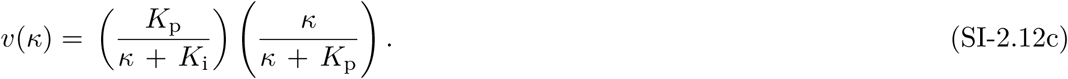

Reference values of parameters used in the expressions (SI-2.11) - (SI-2.12) are listed in Table SI-2.1. These values correspond to the values used to compute Fig. 3 in (Lever et al., 2016).

**Table SI-2.1:** KPL-IFFL model parameter values.

| **№** | **Parameter** | **Value** |
| --- | --- | --- |
| 1. | $K_i$ | $10^{-3}$ |
| 2. | $K_p$ | $10^{-2}$ |
| 3. | $\Gamma^y$ | $5 \times 10^2$ |
| 4. | $\Gamma^p$ | $5 \times 10^2$ |
| 5. | $\Delta$ | $5 \times 10^3$ |
| 6. | $\Theta$ | $5 \times 10^4$ |
| 7. | $\Lambda$ | $10^4$ |
| 8. | $\alpha$ | $10^{-4} - 10^4$ |
| 9. | $\kappa$ | $10^{-4} - 10^2$ |

#### SI-3 A core mathematical model of PD-1 expression

##### SI-3.1 The model equations

Our core mathematical model of PD-1 expression on the surface of a CD8+ T cell, called the KPL-IFFL-PD1 model, describes normal and aberrant dynamics of interactions between four immunobiochemical entities, Bcl-6 (*C*), PD-1 (*P*), IRF4 (*I*), and Blimp-1 (*B*),

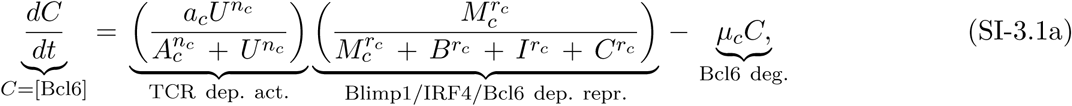

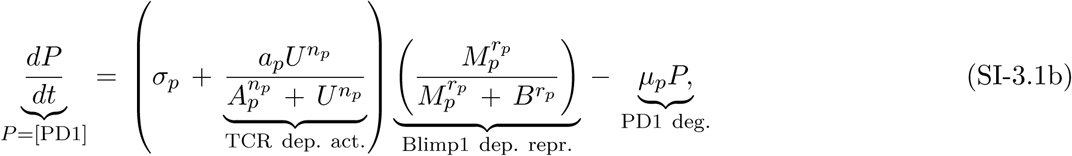

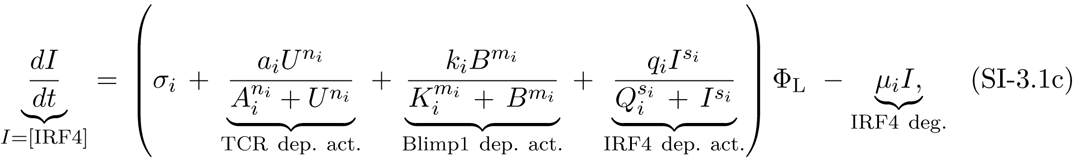

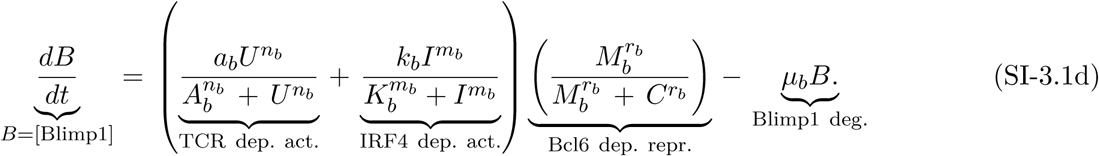

Here, for the sake of compactness in the equation term explanation, we use the following abbreviations, “TCR dep. act.” for TCR-dependent activation, “Blimp-1/IRF4/Bcl-6” for Blimp-1/IRF4/Bcl-6-dependent repression, and so on.

The model structure corresponds to the circuit topology depicted in Fig. 2 with a few simplifications resulting from lumping some species, (*i*) NFATc1 and PD-1 becoming the species *P*, and (*ii*) NF-ϰB and IRF4 becoming the species *I*. We also omit Erk-dependent degradation of Bcl-6 because it is in turn attenuated by Bcl-6 itself.

The input *U* := *U* (*α*, *κ*, *P*) to the KPL-IFFL-PD1 model (SI-3.1) is described by the scalar function *u*(*α*, *κ*) defined in (SI-2.11),

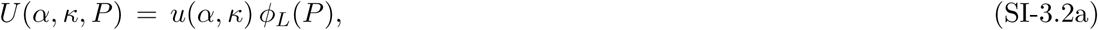

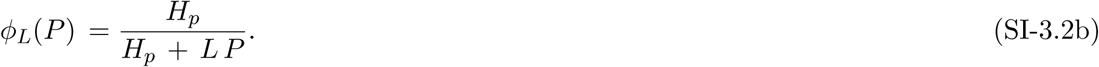

Here, the inhibitory regulatory factor *Φ*_*L*_(*P*) corresponds to the co-localization of PD-1:PD-L1 complexes around the immunologic Ag-TCR synapses that hinder the TCR activity as discussed in Sec. 3.2. An external environment parameter *L* models a fraction of PD-1 receptors bound with PD-L1. Parameters *α* and *κ* are scaled Ag level and scaled *k*_off_, the dissociation constant for the Ag-TLR bond, respectively.

We use a Michaelis-Menton saturation functional dependence in the expression (SI-3.2b) to describe a 2D-sliding diffusion of PD-1 receptors on the surface of a T cell (without any switch-like reaction sharp transitions) as a major process contributing to the TCR down regulation effect (Sect. 3.2).

Next, the factor Φ_L_ := Φ_L_(*P*) in the equation (SI-3.1c) describes a net negative feedback effect caused by the PD-1:PD-L1 interaction (Sec. 3.2),

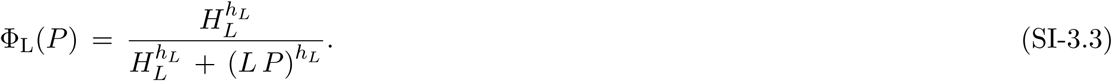

Recall that the active complex formed between PD-1 and PD-L1 suppresses the NF-*κ*B pathway (Sect. 3.4.2), while the NF-*κ*B pathway activates IRF4 (see above).

We use generic Hill functions in (SI-3.2b) and (SI-3.3) following the Hill-function approximation suggested for T cell exhaustion in (Johnson et al., 2011).

In order to capture effects caused by self (tumor) and non-self (infection) interactions, including significant differences in the magnitude of infection and amount of tumor antigens, we implement the following relationships to mathematically implement the self- / non-self specificity,

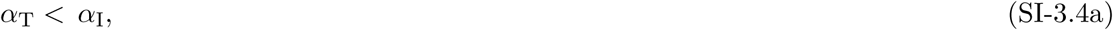

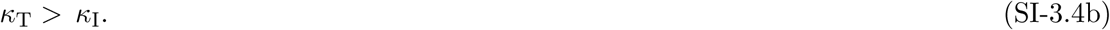

Here, subscript labels “T” and “I” correspond to tumor and infection, respectively.

Based on our immunobiochemical reconstruction and following (Warmflash and Dinner, 2009), we make explicitly additional choices to rank TCR-mediated activation parameters as follows,

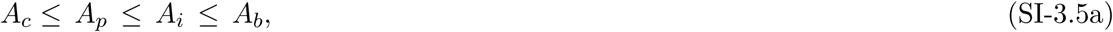

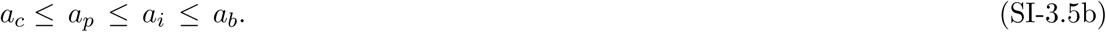

Based on the developed immunobiochemical reconstruction, the inequality choices (SI-3.5a) ensure that the genes encoding Bcl-6 and PD-1 are activated at lower antigen levels than the genes encoding IRF4 and Blimp-1, whereas the latter ensures that the switch towards the suppression of PD-1 transcription is biased towards the CD8+ T cell, when both IRF4 and Blimp-1 are expressed at high antigen levels.

To account for the abundance of the lumped TNF*α*/IFN*γ* species, we have replaced the rate constant *σ*_*p*_ in the equation (SI-3.1b) by the reaction rate expression,

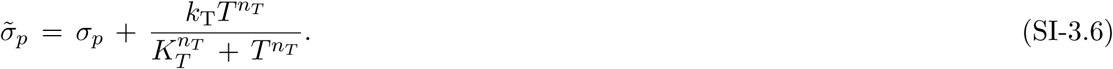

Here, *T* corresponds to TNF*α*, *k*_*T*_ = 0.5, *k*_*T*_ = 1, and *n*_*T*_ = 2. The values of the new parameters are selected in the range of the corresponding parameter values from Table SI-3.1.

##### SI-3.2 The model parameters

Reference parameter values used in our modeling studies are listed in Table SI-3.1. We have to mention explicitly that the parameter values have not been fitted to any data from (Kohlhapp et al., 2016), and have been selected as follows.

**Table SI-3.1:** PD-1 expression model parameter values.

| Parameters | Values | Comments |
| --- | --- | --- |
| $\sigma_i$ | 0.30 | IRF4 constituent synthesis rate |
| $\sigma_p$ | 0.10 | PD-1 immune central tolerance const. synth. rate |
| $a_c = a_p < a_i < a_b$ | $0.75 = 0.75 < 75.0 < 100.00$ | genetic switch thresholds |
| $A_c < A_p < A_i < A_b$ | $0.01 < 0.10 < 1.00 < 10.00$ | genetic switch thresholds |
| $n_c = n_p$ | 3 | species: Bcl-6 and PD-1 |
| $n_b = n_i$ | 2 | species: Blimp-1 and IRF4 |
| $k_b$ | 0 – 25 | species: IRF4 |
| $k_i = q_i$ | 7.50 | species: Blimp-1 and IRF4 |
| $K_b = K_i = Q_i$ | 1.00 | species: Blimp-1 and IRF4 |
| $m_b = m_i = s_i$ | 2 | species: Blimp-1 and IRF4 |
| $M_b = M_c = M_p$ | 10.00 | species: Blimp-1, Bcl-6 and PD-1 |
| $H_p = H_L$ | 0.1 | species: PD-1 and PD-L1 |
| $r_b = r_c$ | 2 | species: Blimp-1 and Bcl-6 |
| $r_p = h_L$ | 4 | species: PD-1 (p) and PD-L1 (L) |
| $\mu_c = \mu_p$ | 0.10 | species: Bcl-6 and PD-1 |
| $\mu_b = \mu_i$ | 1.00 | species: Blimp-1 and IRF4 |
| $L$ | 0 - 1 | species: fraction of PD-1 bound to PD-L1 |

First, we used the same dimensionless (scaled) parameter values for the subset of interactions as those which were used in (Sciammas et al., 2011; Lever et al., 2016).

In our selection of the reference parameter values (Table SI-3.1), we also analyzed and followed a number of insightful discussions of a very challenging and complex problem of selecting relevant parameter values for biological and especially immunological models, presented in a number of published works (Heinrich and Rapoport, 2005; Warmflash and Dinner, 2009; Martinez et al., 2012; Lever et al., 2014; Galvez et al., 2016), including conceptual views (Gunawardena, 2014; Eftimie et al., 2016) as well as discussed general issues with experimental measurements (De Boer and Perelson, 2013; Eftimie et al., 2016).

Second, the parameter values used from (Sciammas et al., 2011; Lever et al., 2016) can be justified for our modeling studies by employing the following IFFL function argument. Indeed, the incoherent feedforward loops (Fig. 2) cannot exert their biphasic function with any arbitrary parameter values (Kim et al., 2008). The parameter values taken from (Sciammas et al., 2011; Lever et al., 2016) and used in the model (SI-3.1) correspond to the dose-dependent biphasic behaviors as defined and studied in (Kim et al., 2008), and also observed experimentally in the cited literature. In other words, the used parameter values are sufficient to instill the IFFL function.

Finally, the type of modeling carried out in our work can be characterized as phenotypic modeling (Warmflash and Dinner, 2009; Lever et al., 2014; Gunawardena, 2014). Recall that the objective of the phenotypic modeling is to capture the function of a biological system, based on the available and well-established features of the regulatory network under study as also explicitly stated in (Sciammas et al., 2011) which justified the selection of generic Hill functions in their model tailored to the GRN topology. In this work, we implemented a similar approach.

#### SI-4 KPL-IFFL-PD1 model additional plots

##### SI-4.1 Master and adaptive roles of the TCR and IRF4

**Figure SI-4.1:**
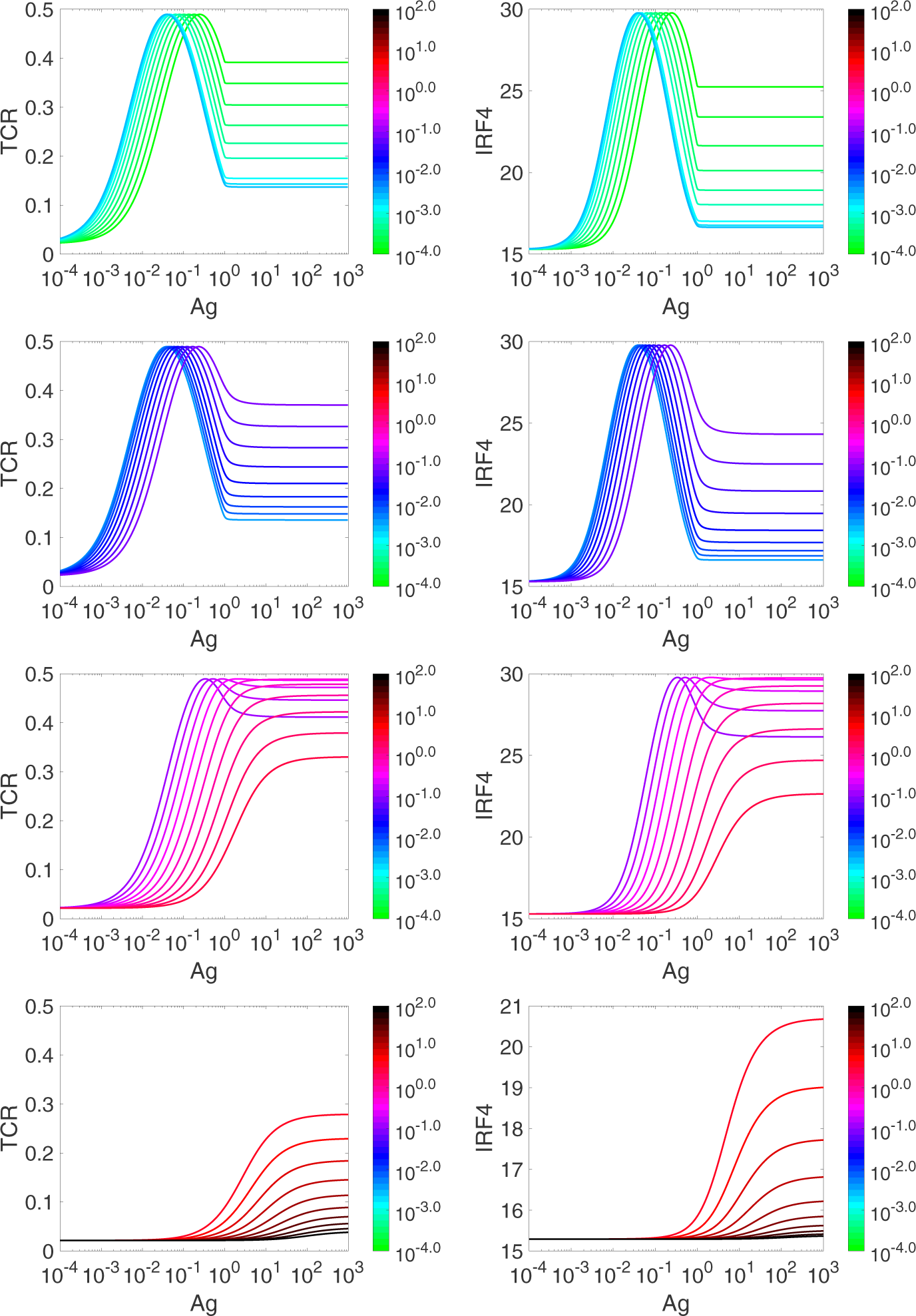
Ag dose-dependent master and adaptive roles of TCR and IRF4, respectively. To generate the shown dependencies, 40 different values for *k*_off_, *k*_off_ ∈ [10^-4^, *-*10^2^], when selected uniformly along the logarithmic scale, log_10_ *k*_off_ ∈ [-4, 2]. The top row corresponds to the first 10 values of the scale, the second row corresponds to the next 10 values, *etc.* The colorbar maps the color-coded plots to the corresponding values of *k*_off_, *k*_off_ = 10^-4^ 10^2^. The panels in the two top rows correspond to the values of *k*_eff_, when the case of “Chronic infection” (Fig. 4) is possible, while the two bottom rows correspond to the case when “Autoimmunity” (Fig. 4) is possible.

##### SI-4.2 Adaptive roles of Bcl-6, Blimp-1, and PD-1 (when IRF4 is on)

**Figure SI-4.2:**
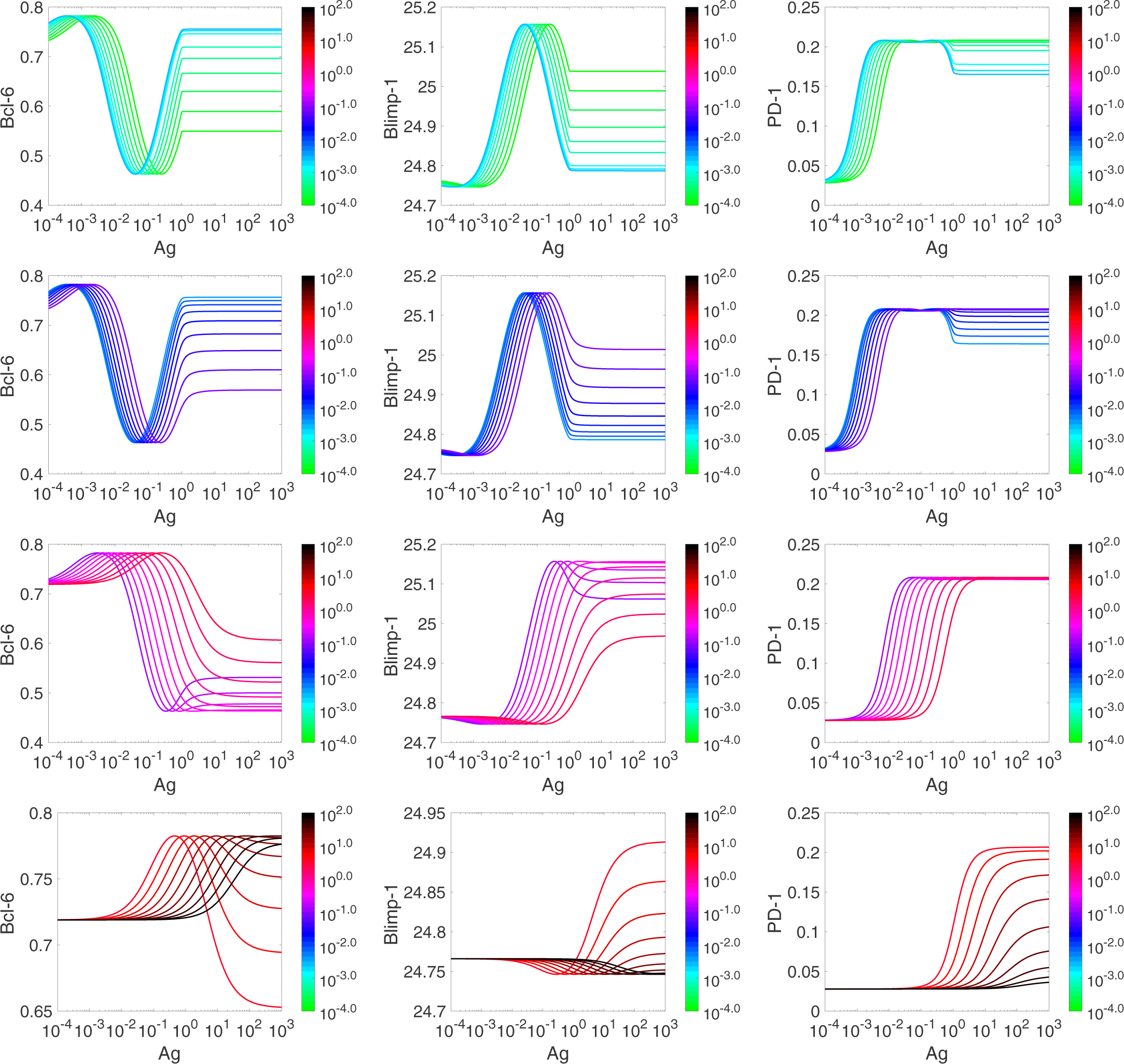
Ag dose-dependent adaptation of Bcl-6, Blimp-1, and PD-1 at different values of *k*_off_, and when IRF4 is turned on. The computations are done as described in the legend for Fig. SI-4.1. The panels in the two top rows correspond to the values of *k*_eff_, when the case of “Chronic infection” (Fig. 4) is possible, while the two bottom rows correspond to the case when “Autoimmunity” (Fig. 4) is possible.

##### SI-4.3 Adaptive roles of Bcl-6, Blimp-1, and PD-1 (when IRF4 is off)

**Figure SI-4.3:**
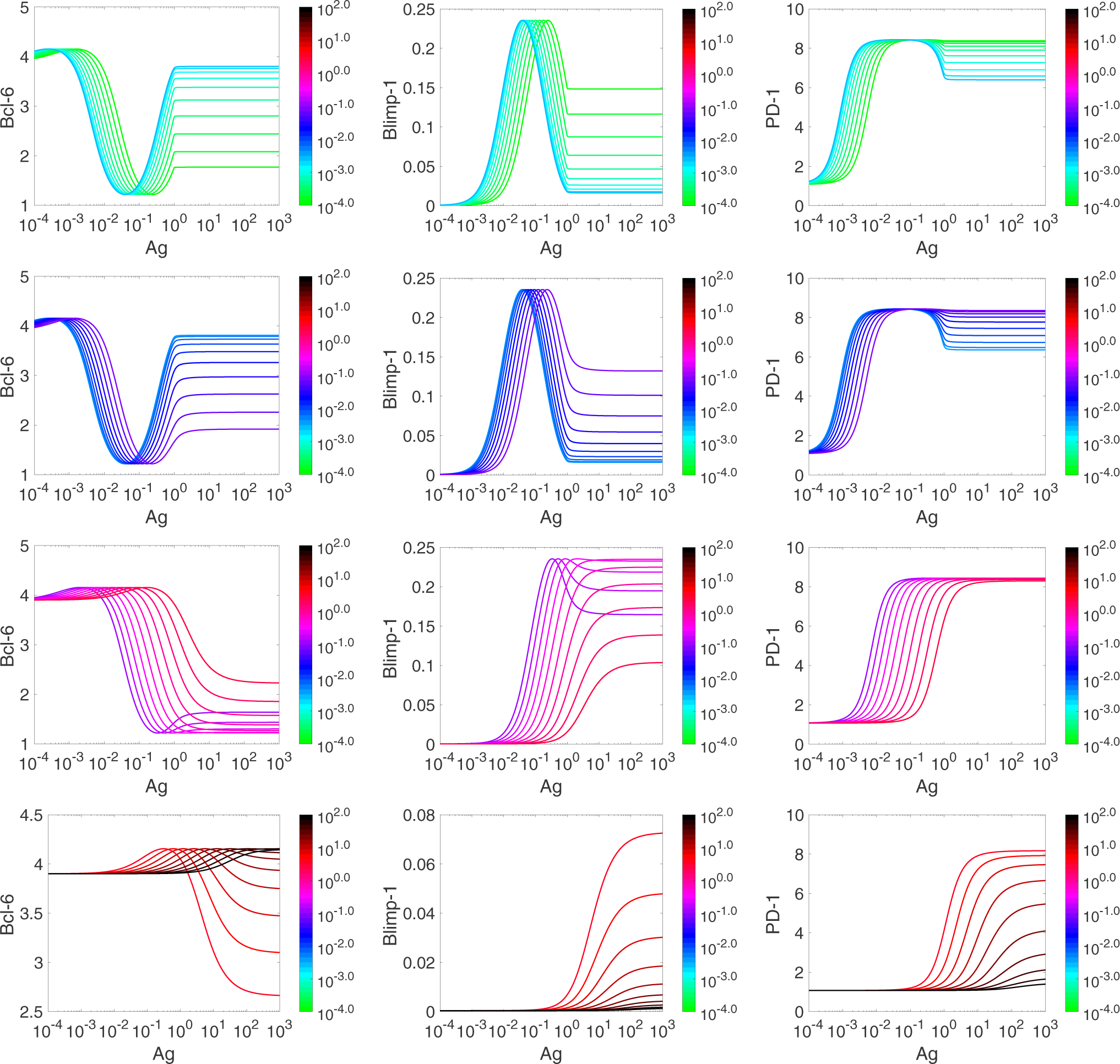
Ag dose-dependent adaptation of Bcl-6, Blimp-1, and PD-1 at different values of *k*_off_ and IRF4 turned off. All computations are done as those to generate Fig. SI-4.2 with one exception that the zero value of *k*_*b*_ (*k*_*b*_ = 0) in the equation (SI-3.1d) to disconnect the rheostat IRF4 from the bottom part of the GRN depicted in Fig. 2. The panels in the two top rows correspond to the values of *k*_eff_, when the case of “Chronic infection” (Fig. 4) is possible, while the two bottom rows correspond to the case when “Autoimmunity” (Fig. 4) is possible.

##### SI-4.4 Modeling PD-1:PD-L1 interactions

###### SI-4.4.1 TCR activation plots

**Figure SI-4.4:**
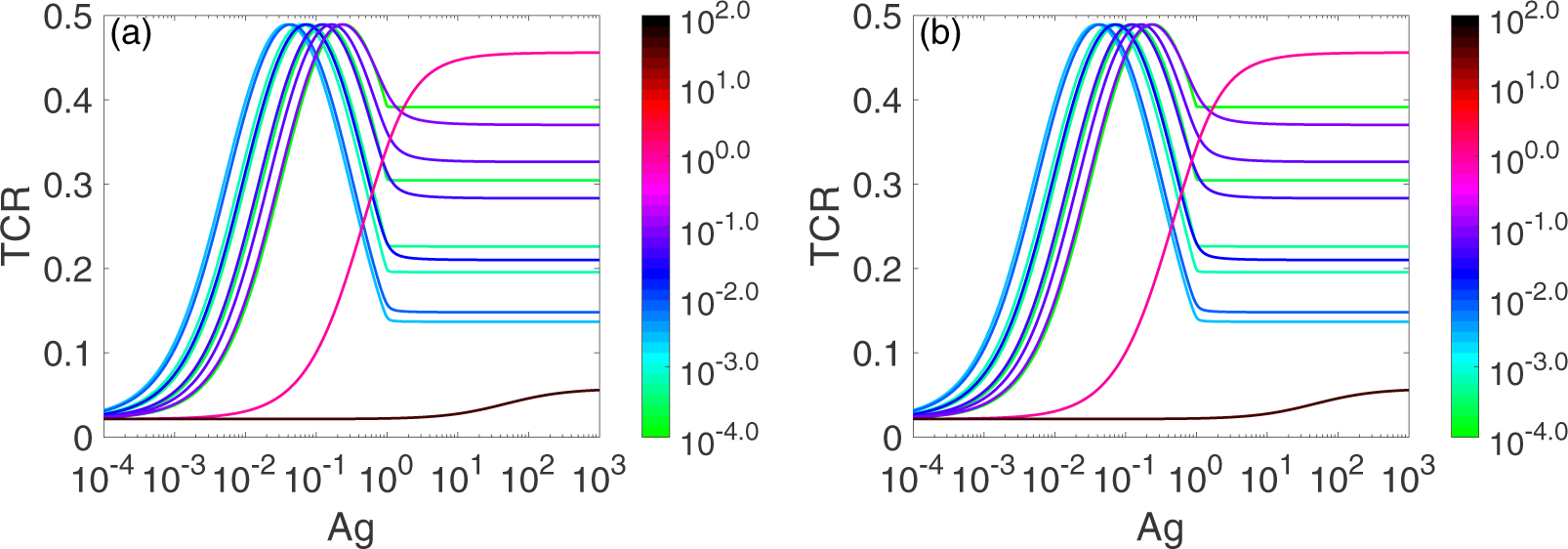
Ag dose-dependent TCR responses at different values of *L* and *k*_eff_. Panels (a), (b), and (c) correspond to *L* = 0.15, 0.5, and 0.9. Colors in the colorbar encode the corresponding 12 values of the parameter *k*_off_ used in the simulations, *k*_off_ *∈ {*10^-4^, 2.03 × 10^-4^, 4.13 × 10^-4^, 5.88 × 10^-4^, 2.43 × 10^-3^, 7.01 × 10^-3^, 2.03 × 10^-2^, 4.13 × 10^-2^, 5.88 × 10^-2^, 8.38 × 10^-2^, 1.0, 49.24*}*.

###### SI-4.4.2 Bcl-6, IRF4, Blimp-1, and PD-1 level plots

**Figure SI-4.5:**
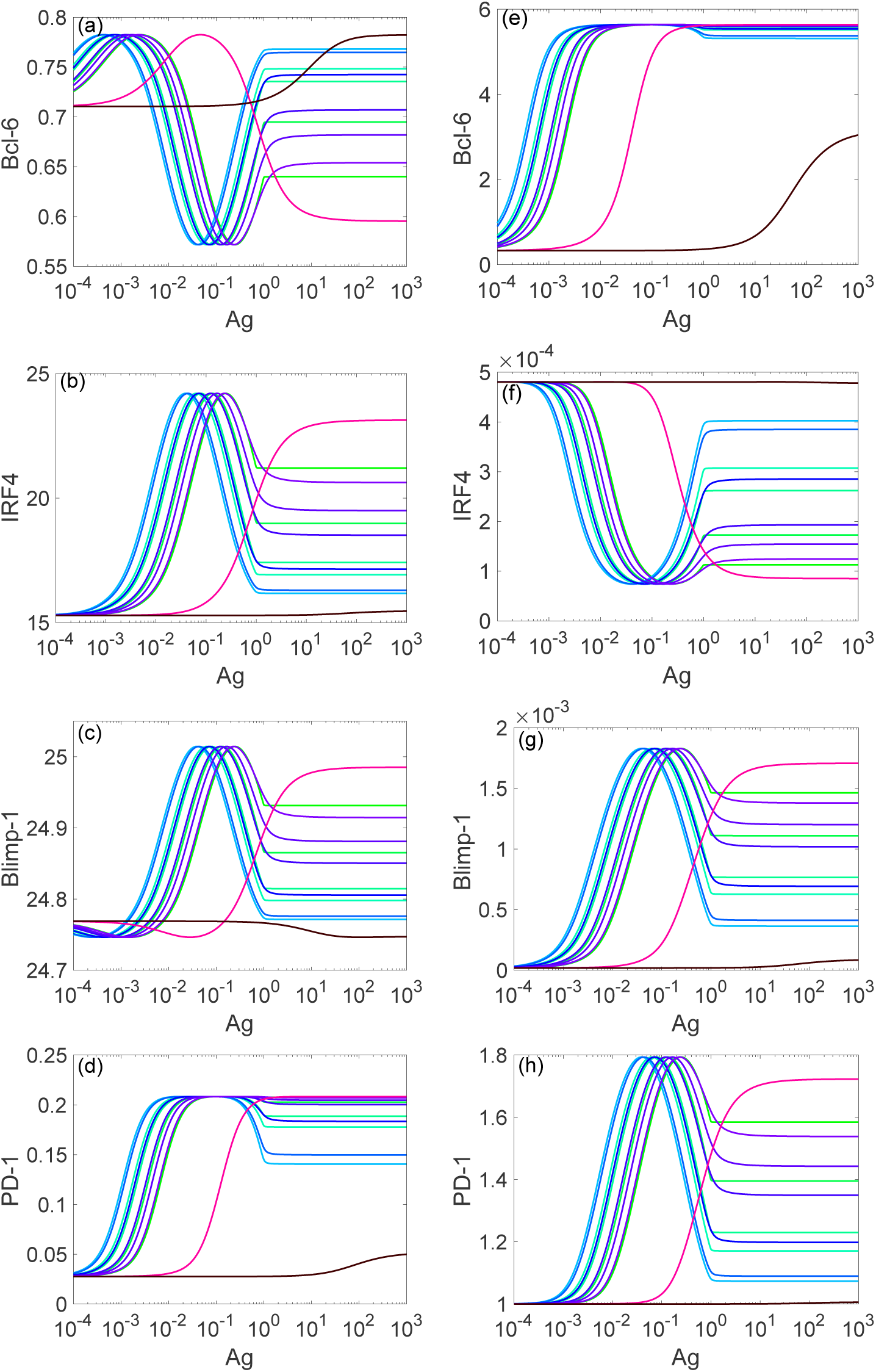
Ag-dose dependent responses of Bcl-6, IRF4, Blimp-1, and PD-1 at different values of *L* and *k*_eff_. All computations are done as described in the legend of Fig. SI-4.4 Panels (a) - correspond to *L* = 0.15, while panes (e) - (h) correspond to *L* = 0.5.

#### SI-5 Estimation of the impact of cell-to-cell variability on TCR activation and PD-1 expression

To capture the heterogeneity of T cell activation (Sect. 3.5.1) more systematically, we employed Global Sensitivity Analysis (GSA) (Saltelli et al., 2008; Ghanem et al., 2017) developed to assess how uncertainty in model parameters propagates and affects the model output, which is in our case either (*i*) the strength of the TCR activation (Fig. 4) or (*ii*) the level of PD-1 expressed on the surface of a T cell (Figs. 6 and 7, Panel (a)).

Technologically, the GSA can provide reliable pre-experimental estimates on the sensitivities of the TIFFL-model properties with respect to broad scale variations in the TIFFL model parameters without knowing their precise values, which is exactly the case in our modeling studies (Sect. SI-3.2). We use the theoretical results obtained using the GSA to provide useful insights into key experiential observations (Kohlhapp et al., 2016).

##### SI-5.1 Case I. The GSA of the TCR activation function

In the case of the TCR activation model (Sect. 3.5.1), we analyzed the level of TCR activation potential to induce downstream signaling pathways (Lever et al., 2016), which is mathematically represented in terms of the values of the function *u*(*α, κ,* Θ, Λ, Δ) defined in (SI-2.11).

Recall that *α* is the scaled concentration of antigen ([Ag]); *κ* is the scaled rate constant *k*_off_, the off-rate constant for Ag and TCR; Θ is the scaled rate constant *µ*, describing repression of the TCR activation within the immunological synapse (Fig. SI-2.1); Λ is the scaled rate constant *λ*, describing amplification of the TCR activation within the immunological synapse (Fig. SI-2.1); and Δ is the scaled rate constant *δ*, describing activation of the TCR within the immunological synapse (Fig. SI-2.1). All these parameters characterize the properties of the corresponding IFFL (Fig. SI-2.1) as defined in (Lever et al., 2016).

Recall also that a variety of ligand-receptor interactions take place at the immunological synapse, many of which are inhibitory (Sect. 3.2), and the final integration between activatory and inhibitory interactions sets the degree of T cell activation. In this respect, we interpret the five model parameters as quantifying a myriad of complex processes that occur at the immunological synapse.

Assuming that uncertainty in the values of the five parameters can be attributed to both environment and cell-to-cell variability, we use the GSA to estimate to which extent the uncertainty and variation in the values of each parameter can contribute to the variation in the strength of TCR activation of the corresponding downstream signaling pathways.

Let the total variation or uncertainty in the values of the TCR activation function *u*(*α, κ,* Θ, Λ, Δ) constitute 100%, or one if normalized and fractions are used. In order to rank factors that contribute to the TCR activation uncertainty originating from both cell-to-cell and environment variability, one can use global sensitivity indexes (Table SI-5.1), computed over the entire set of all parameter values as discussed in Sect. SI-6.

**Table SI-5.1:** Sobol’s first-order and total sensitivity indexes for *u*(*α*, *κ*, Θ, Λ, Δ).

| **Nº** | **Factor<sup>a</sup>** | **Single Effect**<br>(First-order index $S_i$) | **Total Effect**<br>(Total index $S_{Ti}$) | **Function** |
| --- | --- | --- | --- | --- |
| 1. | $\Delta$ | 0.282 | 0.477 | TCR activation ($\delta$) |
| 2. | $\kappa$ | 0.115 | 0.366 | Ag-TCR affinity ($k_{\text{off}}$) |
| 3. | $\alpha$ | 0.112 | 0.350 | Ag level ($[\text{Ag}]$) |
| 4. | $\Theta$ | 0.059 | 0.165 | TCR inhibition ($\mu$) |
| 5. | $\Lambda$ | 0.057 | 0.147 | TCR amplification ($\lambda$) |

<sup>a</sup> In the GSA, parameters are called factors, $i \in \{\Delta, \kappa, \alpha, \Theta, \Lambda\}$ .

In Table SI-5.1, factors 1, 2, 4, and 5 correspond to cell-to-cell variability, while factor 2 corresponds to tissue characteristics variability (Ag-levels).

Recall that the first-order sensitivity index *S*_*i*_ is computed in two steps (Saltelli et al., 2008). First, the mean of the output function *u* is computed conditioned to the factor *F*_*i*_ (or, equivalently, the *i*-th model parameter used in the analysis), and so, the mean is a function of its argument 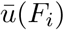. Then, the variance of 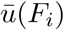 with respect to all allowable values of *F*_*i*_ is calculated and normalized to the total variance in the values of *u* with respect to changes in all factors (parameters). If the function 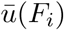 is not a constant function of its argument, this will lead to non-zero variance and, as a result, a non-zero value of the index *S*_*i*_. The computation of the total factor reveals non-linear interactions between the given factor *F*_*i*_ and all other factors and we omit the details (Saltelli et al., 2008).

From the analysis of Table SI-5.1, we can conclude that three parameters, Δ, *κ*, and *α*, can be ranked as the most influential parameters. Assuming that evolution has shaped the immune response towards the clearance of (non-self) infection as an “ideal” primary function of immunity, the loss of the functional response to tumor self-antigens can constitute about 50.87% due to the corresponding variations in the TCR activation within the immunologic synapse (∼28.15%), the TCR “affinity” (∼11.52%), and the antigen level (∼11.20%).

In this respect, we can interpret the larger total effects (Table SI-5.1) as amplification of the corresponding single parameter variability impacts due to the interactions with other factors.

This interpretation and quantification of uncertainty should be used with care as a myriad of other factors as well as the interactions among the factors are not considered. However, due to the well-established practical use of the GSA and the normalization condition used in the GSA, when the sum of all *positive* sensitivity indexes must add to one, only a handful of sensitivity coefficients can be large, while the majority of the sensitivity coefficients should be negligibly small (Saltelli et al., 2008). This means that irrespectively of the corresponding numerical values, the most influential factors are likely to be identified correctly.

Note that the discussed situation in the GSA is somewhat similar to what is also very well known in the MCA (Metabolic Control Analysis) (Kacser and Burns, 1973; Heinrich and Rapoport, 1974), which correctly identifies a handful of bottleneck enzymes in the given metabolic pathway at the given (local) reference state, while the majority of the enzymes operate under equilibrium condition and, thus, cannot influence the flux through the pathway significantly (Heinrich and Schuster, 1996; Nikolaev, 2010).

##### SI-5.2 Case II. The GSA of the PD-1 steady-state level in the presence of PD-1:PD-L1 interaction

We next estimate Sobol’s sensitivity indexes (Table SI-5.2) for the model that includes the interaction between PD-1 and PD-L1 (Sect. 3.5.3).

To check the robustness of the GSA-based factor ranking, we have computed the Sobol’s indexes for the two sets of factors included in the model analysis (Table SI-5.2).

**Table SI-5.2:** Sobol’s first-order and total sensitivity indexes for PD-1 steady-state levels in the presence of the interaction PD-1:PD-L1.

| **Nº** | **Factor** | **Single Effect**<br>(First-order index $S_i$) | **Total Effect**<br>(Total index $S_{Ti}$) | **Function** |
| --- | --- | --- | --- | --- |
| 1. | $L$ | 0.320 (0.528)<sup>a</sup> | 0.615 (0.775) | PD-1:PD-L1 interaction ($L$) |
| 2. | $\Delta$ | 0.104 (0.090) | 0.363 (0.356) | TCR activation ($\delta$) |
| 3. | $\kappa$ | 0.043 (0.020) | 0.283 (0.234) | Ag-TCR affinity ($k_{\text{off}}$) |
| 4. | $\alpha$ | 0.036 (0.007) | 0.249 (0.207) | Ag level ($[\text{Ag}]$) |
| 5. | $\Theta$ | 0.023 | 0.133 | TCR inhibition ($\mu$) |
| 6. | $\Lambda$ | 0.019 | 0.117 | TCR amplification ($\lambda$) |

<sup>a</sup> Indexes shown in parentheses are computed for the fixed reference values of the parameters $\Theta$ and $\Lambda$ (Table SI-2.1), called “Factor Prioritization” in the GSA; $i \in \{L, \Delta, \kappa, \alpha, \Theta, \Lambda\}$ .

We observe from Table SI-5.2 that the two most influential factors leading to the high variability in the steady-state levels of the PD-1 state variable include two parameters, (*i*) *L*, the fraction of PD-1 receptors ligated with PD-L1, and (*ii*) Δ, the scaled rate constant corresponding to the activation of the TCR in the immunologic synapse.

The data show that the index *S*_*L*_ stands out of all other indexes, confirming that this environment factor plays a significant role in the expression of PD-1.

At the same time, the fact that both indexes *S*_*L*_ and *S*_Δ_ have the largest values additionally emphasizes the duality of PD-1 used as a biomarker of both activated and exausted CD8+ T cells as discussed earlier in Mechanism **(O1-M6)** of Sect. 3.3.

##### SI-5.3 Case III. The GSA of the PD-1 steady-state level in the presence of PD-1:PD-L1 interaction and pro-inflammatory cytokines TNF*α***/INF***γ*

We now add an additional factor *T* into the model (SI-3.1), using the updated rate expression (SI-3.6). The factor *T* describes the scaled level of lumped proinflammatory cytokines TNF*α* and INF*γ*. For the updated model, the corresponding Sobol’s indexes are collected in Table SI-5.3

**Table SI-5.3:** Sobol’s first-order and total sensitivity indexes for PD-1 steady-state levels in the presence of the interaction PD-1:PD-L1 and pro-inflammatory cytokines TNF*α*/INF*γ*.

| **Nº** | **Factor** | **Single Effect**<br>(First-order index $S_i$) | **Total Effect**<br>(Total index $S_{Ti}$) | **Function** |
| --- | --- | --- | --- | --- |
| 1. | $L$ | 0.656 | 0.732 | PD-1:PD-L1 interaction ($L$) |
| 2. | $T$ | 0.267 | 0.318 | $\text{TNF}\alpha/\text{INF}\gamma$ ($T$) |
| 3. | $\Delta$ | 0.003 | 0.026 | TCR activation ($\delta$) |
| 4. | $\kappa$ | 0.003 | 0.017 | Ag-TCR affinity ($k_{\text{off}}$) |
| 5. | $\alpha$ | $5.038 \times 10^{-4}$ | 0.017 | Ag level ($[\text{Ag}]$) |

We observe from Table SI-5.3 that the most influential factors related to the variations in the PD-1 levels correspond to (*i*) the interaction PD-1:PD-L1 and (*ii*) the inflammation state of the environment.

From our comparison of the results in Tables SI-5.2 and SI-5.3, we can conclude that the factor corresponding to the PD-1:PD-L1 interaction is amplified in the presence of the pro-inflammatory cytokines.

The experimental results (Kohlhapp et al., 2016) show that anti-influenza CD8+ T cells have much smaller numbers of PD-1 receptors expressed on their surface as compared with the anti-melanoma CD8+ T cells shunted to the infected lung. This means that due to the largest value of the sensitivity index *S*_*L*_, the processes leading to the down-regulation of PD-1 transcription via the discussed TIFFL mechanism should not be significantly affected in anti-influenza CD8+ T cells.

Indeed, the “sensitivity” of the corresponding processes works in both directions: (*i*) when the intensity of the PD-1:PD-L1 interaction rapidly increases, the level of PD-1 should also rapidly change, in our case, it should increase, and vise versa, (*ii*) as soon as the intensity of the PD-1:PD-L1 interaction rapidly decreases, the “brake” from the TIFFL mechanism should also be rapidly removed due to the quantified sensitivity of the mechanism to the PD-1:PD-L1 interaction.

##### SI-5.4 Case IV. The GSA of the PD-1 steady-state level in the presence of PD-1:PD-L1 interaction, pro-inflammatory cytokines TNF*α***/INF***γ*, and IL-2

We finally discuss the Sobol’s sensitivity indexes due to the variation in the parameter *γ* used in the scaling product *γ a*_*b*_, leading to the increase in the level of Blimp-1 caused by the large amounts of IL-2 as discussed in Sect. 3.5.4.

In this case, our simulations have revealed that the changes in the value of the scaling factor *γ* ≥ 1 affect the level of PD-1 in a stepwise fashion in the model (SI-3.1). This can be attributed to the current coarse-grained structure of the model to capture the IL-2 effect: There is a threshold *γ*^0^ at which the steady-state level of PD-1 drops sharply to almost zero values for *γ* > *γ*^0^, while it is not affected for *γ* < *γ*^0^. This results in the almost constant value of the mean of 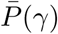 conditioned to *γ* and computed with respect to all other parameter values, leading to the lower value of the corresponding first order sensitivity index *S*_*γ*_ (Table SI-5.4).

**Table SI-5.4:** Sobol’s sensitivity indexes for PD-1 steady-state levels in the presence of (*i*) the interaction PD-1:PD-L1, (*ii*) TNF*α*/INF*γ*, and (*iii*) IL-2.

| Nº | Factor | Single Effect<br>(First-order index $S_i$) | Total Effect<br>(Total index $S_{Ti}$) | Function |
| --- | --- | --- | --- | --- |
| 1. | $L$ | 0.517 | 0.651 | PD-1:PD-L1 interaction ($L$) |
| 2. | $T$ | 0.200 | 0.246 | $\text{TNF}\alpha/\text{INF}\gamma$ ($T$) |
| 3. | $\gamma$ | 0.037 | 0.216 | Blimp-2 transcription |
| 4. | $\Delta$ | 0.010 | 0.162 | TCR activation ($\delta$) |
| 5. | $\kappa$ | 0.001 | 0.110 | Ag-TCR affinity ($k_{\text{off}}$) |
| 6. | $\alpha$ | 0.006 | 0.098 | Ag level ($[\text{Ag}]$) |

However, the total effect index *S*_*T* *γ*_ computed for the factor *γ* turns out to be large, meaning that the factor of IL-2 strongly interacts with other factors in the model (Table SI-5.4).

Our modeling results further point out to interesting trends in the Sobol’s sensitivity indexes (Table SI-5.4). Specifically, the IL-2 factor is ranked as the third most important factor. Moreover, its total effect index (*S*_*T* *γ*_ = 0.216) is comparable with the total effect index for TNF*α*/INF*γ* (*S*_*T*_*T* = 0.246). As such, the total effect induced by IL-2 has a potential to counteract the total effect caused by TNF*α*/INF*γ*.

Besides, while the indexes corresponding to the PD-1:PD-L1 and TNF*α*/INF*γ* factors are decreased (cp., Tables SI-5.3 and SI-5.4), the total effect indexes corresponding to both the TCR activation and Ag-related factors are considerably increased, meaning that the abundance of influenza A-infected cells in the lung can positively affect signaling pathways leading to the activation of Blimp-1 expression in anti-melanoma CD8+ T cells, and as a result of the suppression of PD-1 expression (Fig. 3).

We can finally interpret the data of Table SI-5.4 in the light of PD-1 blockade experimental results (Kohlhapp et al., 2016) as removing the most influential PD-1:PD-L1 factor. In this case, the other two influential TNF*α*/INF*γ* and IL-2 factors counteract one another leading to the activation of both IRF4 and Blimp-1 pathways, which are hallmarks of reactivated (anti-melanoma) CD8+ T cells as discussed around Fig. 3.

#### SI-6 GSA computational method

We used two types of Sobol’s sensitivity indexes, including (Saltelli et al., 2008; Cannavó, 2012; Ghanem et al., 2017):

(*i*) first-order indexes, *S*_*i*_, *i* = 1,…, *n*, which take into account the variance of the mean value of the output function, *e.g.*, *u*(*α*, *κ*, Θ, Λ, Δ), depending on all parameters, conditioned to the given factor *F*_*i*_, with respect to all values of the factor, estimated over the given multidimensional parameter hypercube. Here, the conditioned mean of the output function is the best predictor of the function based on *F*_*i*_.

(*ii*) total effect indexes, *S*_*T* *i*_, *i* = 1,…, *n* which take into account interaction of the given factor with all other factors used in the analysis; here, for example, if *n* = 3 then *S*_*T* 1_ = *S*_1_ + *S*_12_ + *S*_13_ + *S*_123_ includes all terms capturing factor *F*_1_ and its interactions with other two factors, *F*_2_ and *F*_3_.

In order to define the multidimensional hypercube used in Monte-Carlos sampling, the given ranges for the parameter values were used if known *a priori* (Fig. 4) or the parameter value lower and upper bounds were estimated using the given reference parameter value as follows. Let *p*_0_0be a reference parameter value as given in Tables SI-3.1 or SI-2.1, then we set 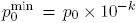 and 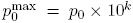. In other words, in all such cases we allow the parameter value to vary within two or four orders of magnitude, the approach corresponding to a recommended parameter estimation practice suggested and used in (Aldridge et al., 2006; Eydgahi et al., 2013).

In the case where the output faction used in the GSA corresponds to a state variable evaluated at steady-state (*e.g.*, *P* = PD-1_*ss*_), for each sample of parameter values the corresponding model was integrated with the same unit initial conditions over the time interval *t* = 300 until the solution reaches its steady-state numerically, using the Matlab^®^-based function ode45.

Given *n* model parameters used in the GSA, the numerical accuracy of the computed Sobol’s sensitivity indexes (Saltelli et al., 2008) were tested in all computationally feasible cases by computing all of the 2^*n*^ - 1 indexes (accounting for both single and group-interaction effects), followed by summing up the computed 2^*n*^ - 1 indexes to obtain the sum equal to one (Saltelli et al., 2008; Cannavó, 2012).

